# Multicore fiber optic imaging reveals that astrocyte calcium activity in the cerebral cortex is modulated by internal motivational state

**DOI:** 10.1101/2023.05.18.541390

**Authors:** Yung-Tian A. Gau, Eric Hsu, Jaepyeong Cha, Rebecca W. Pak, Loren L. Looger, Jin U. Kang, Dwight E. Bergles

## Abstract

Astrocytes are a direct target of neuromodulators and can influence neuronal activity on broad spatial and temporal scales through their close proximity to synapses. However, our knowledge about how astrocytes are functionally recruited during different animal behaviors and their diverse effects on the CNS remains limited. To enable measurement of astrocyte activity patterns *in vivo* during normative behaviors, we developed a high-resolution, long working distance, multi-core fiber optic imaging platform that allows visualization of cortical astrocyte calcium transients through a cranial window in freely moving mice. Using this platform, we defined the spatiotemporal dynamics of astrocytes during diverse behaviors, ranging from circadian fluctuations to novelty exploration, showing that astrocyte activity patterns are more variable and less synchronous than apparent in head-immobilized imaging conditions. Although the activity of astrocytes in visual cortex was highly synchronized during quiescence to arousal transitions, individual astrocytes often exhibited distinct thresholds and activity patterns during explorative behaviors, in accordance with their molecular diversity, allowing temporal sequencing across the astrocyte network. Imaging astrocyte activity during self-initiated behaviors revealed that noradrenergic and cholinergic systems act synergistically to recruit astrocytes during state transitions associated with arousal and attention, which was profoundly modulated by internal state. The distinct activity patterns exhibited by astrocytes in the cerebral cortex may provide a means to vary their neuromodulatory influence in response to different behaviors and internal states.

## Introduction

Astrocytes are ubiquitous and essential components of neural circuits in the mammalian CNS that are organized in a grid-like manner in the parenchyma, with each cell extending highly ramified processes to allow extensive interactions with neurons, glial cells, and the vasculature (Barber et al., 2021). The fine lamellar protrusions that project from their processes separate and often encircle synapses, creating a “tripartite” structure with pre- and postsynaptic elements (Papouin et al., 2017a). It has been estimated that each astrocyte contacts an enormous number of synapses (about 90,000 in rodents and 2,000,000 in humans) (Oberheim et al., 2012; Refaeli et al., 2021), allowing each cell to serve as a hub to integrate input from diverse sources to modulate the physiological output of neurons on broad spatial and temporal scales (Ackerman et al., 2021; Chaturvedi et al., 2022; Doron et al., 2022; Hsu et al., 2021; Kol et al., 2020; Mu et al., 2019; Nagai et al., 2021; Yu et al., 2020). This functional integration of astrocytes is enabled in part by their expression of a diverse array of metabotropic receptors that can increase intracellular levels of calcium and other second messengers, triggering pleotropic effects ranging from regulation of glycolysis (Bélanger and Magistretti, 2022) to release of “gliotransmitters” that can influence neuronal activity patterns and synaptic strength (Haydon, 2017; Requie et al., 2022; Yu *et al*., 2020). Recent single-cell mRNA sequencing studies have revealed that individual astrocytes exhibit distinct molecular characteristics and that translation occurs in their highly ramified processes, creating a framework for regionally-specific responses to global neuromodulation (Akdemir et al., 2022; Bayraktar et al., 2020; Bouvier et al., 2016; Endo et al., 2022; Gingrich et al., 2022; Miller et al., 2019; Sakers et al., 2021; Sapkota et al., 2022). Although functional validation of this regional specification has started to emerge, many questions remain to be answered regarding the spatial extent and timing of their activity during different behaviors (Maloney et al., 2022), the mechanisms that control their responses (Sardar et al., 2021), and the consequences of their distinct activity patterns on local circuitry (Ung et al., 2020). Understanding the complex role of astrocytes during different behavioral states has been limited by a paucity of information about the spatial and temporal dynamics of astrocyte activity during distinct voluntary behaviors (Nimmerjahn and Bergles, 2015).

Neuromodulatory systems are vital for our capacity to adapt and adjust behaviors based on internal states and changes in environment. In particular, these systems provide the means to register internal states, such as starvation versus satiety, or somnolence versus wakefulness. They can also convey information about external cues, such as the emergence of nutritional resources or unexpected rewards and novelties to enable adaptive behaviors (D’yakonova, 2014). As astrocytes are a direct target of neurotransmitters, including norepinephrine (NE), dopamine (DA) and acetylcholine (ACh) (Corkrum et al., 2020; Ingiosi and Frank, 2022; Kjaerby et al., 2020; Papouin et al., 2017b; Reitman et al., 2023; Rungta et al., 2016; Srinivasan et al., 2015), and can be coupled into functional units (Szabó et al., 2017), with access to many thousands of synapses, they are well-positioned to act both as an amplifier through which sparse neuromodulatory inputs convey contextual changes, and as a gain modulator to fine-tune local circuit output (Corkrum *et al*., 2020; Doron *et al*., 2022; Farhy-Tselnicker et al., 2021; Kol *et al*., 2020; Refaeli *et al*., 2021; Szabó *et al*., 2017).

Understanding the relationship between the engagement of astrocytes and animal behavior requires the ability to monitor astrocyte activity patterns while animals are engaged in physiologically relevant behaviors (Oliveira et al., 2015). This has now been realized through the development of genetically encoded calcium sensors, cell-specific molecular targeting, and *in vivo* brain imaging. Nonetheless, *in vivo* imaging in the mouse brain commonly requires head immobilization to limit motion artifacts and enable use of heavy and complex imaging objectives. This approach is suitable for delivery of controlled stimuli and stereotyped training paradigms; however, restraining precludes full recapitulation of the complex behaviors that an animal performs in adapting to different contexts (Miyamoto and Murayama, 2016) and may be confounded by heightened arousal and stress. While head-mounted mini-microscopes (Ghosh et al., 2011) have been widely used to visualize neuronal activity in freely moving animals, application of this approach has been more difficult for astrocytes (Ingiosi et al., 2020), with most information about astrocyte calcium dynamics in freely behaving mice obtained using photometry rather than cellular imaging (Qin et al., 2020; Sekiguchi et al., 2016; Zhang et al., 2021). Astrocytes present unique challenges for *in vivo* imaging studies, due to their unique morphological features and physiological properties. In particular, they exhibit dramatic structural and physiological changes to brain injury, a transformation termed reactive astrocytosis (Escartin et al., 2021; Hsu *et al*., 2021; Zamanian et al., 2012), which is induced when micro-lenses used in mini-microscopes are implanted into the brain. This issue is further complicated by the scattering nature of brain tissue (and particularly at scar boundaries), making it difficult to image beyond the layer of reactive astrocytosis (Lee et al., 2016). Moreover, the dense packing and highly ramified, thin processes of astrocytes (Hama et al., 2004) place acute demands on the resolving power of imaging systems (Song et al., 2017). Resolution of fluorescence changes induced by genetically encoded calcium sensors is also more difficult in astrocytes, as their calcium transients arise predominantly through internal store mobilization, which are generally smaller events than those in neurons arising though action potential- dependent gating of calcium channels (Gee et al., 2015; Ho and van den Pol, 2007). Although astrocyte metabotropic receptor-induced calcium transients in astrocytes are more prolonged, their small amplitude necessitates efficient photon capture, which is constrained by the small lenses employed in these devices.

To enable visualization of astrocyte calcium changes within the cerebral cortex of freely moving animals, we developed a widefield, long working distance fiber optic imaging system capable of resolving cellular changes in fluorescence emitted by genetically encoded calcium indicators *in vivo*. Taking advantage of spatial-division multiplexing optics, this approach uses a multi-core optical fiber coupled to miniature lenses that provide sufficient working distance to be mounted outside of a cranial window, minimizing brain injury (Cha and Kang, 2014; Cha et al., 2015; Pak et al., 2021). This platform enables repetitive imaging over long periods to delineate the activity patterns of both individual astrocytes and multicellular astrocyte networks in the upper layers of the cerebral cortex, which receives extensive neuromodulatory input. Using this approach, we defined the spatiotemporal dynamics of astrocytes during diverse behaviors, demonstrated that noradrenergic and cholinergic efferents coordinate to recruit astrocytes during various levels of arousal and attention, and showed that the detection threshold of astrocytes to a stimulus can be profoundly altered by internal state. This novel imaging platform provides a means to better study how regulation of astrocyte physiology by neuromodulators influences activity in discrete circuits to modify information processing and behavior.

## Results

### Spatial-division multiplexed fiber optic imaging allows visualization of astrocyte activity *in vivo*

Astrocytes in the upper layers of the cerebral cortex exhibit robust increases in calcium in response to behaviorally-induced changes in neural activity (Merten et al., 2021; Paukert et al., 2014; Srinivasan *et al*., 2015; Thrane et al., 2012), providing an opportunity to assess astrocyte network activity in freely moving animals using minimally invasive imaging through implanted optical windows. To define the behavioral contexts during which astrocytes are activated, we developed a flexible fiber optic imaging system that provides both the cellular resolution necessary to monitor fluorescence changes from individual astrocytes, and a field of view wide enough to visualize networks of astrocytes in these circuits. Imaging was achieved using a spatial-division multicore optical fiber in which 30,000 individual quasi-single mode glass fiber cores are assembled into a circular array < 1 mm in total diameter with an inter-fiber spacing of 2.1 µm. Flexible multicore fibers have been previously exploited for photometric analysis (Ferezou et al., 2006) and low-resolution, low signal-to-noise ratio detection of fluorescence changes in the brain (Flusberg et al., 2008; Paukert *et al*., 2014). To perform cellular resolution imaging at a distance, so as to avoid penetrating injury and tissue compression that occurs when imaging devices with a short focal distance, such as gradient-index lenses (Salatino et al., 2017), are inserted directly into the brain, we coupled the end of the multicore fiber to two miniature aspheric lens doublets, together serving as the objective lens, achieving a numerical aperture of 0.58, yet retaining a working distance sufficiently long (2.5 mm) to allow imaging through a cranial window (Figure 1A, B). Fluorescence imaging was performed using a 473 nm diode laser as an excitation source and a charge-coupled device (CCD) camera as a detector (see Methods), a configuration that achieved a lateral resolution of 2.8 µm and an axial resolutionof 24 µm (Figure 1C). Although used in an epifluorescence imaging mode, the small 2.1-µm aperture of each fiber enhances optical sectioning by acting as a pinhole to minimize detection of light from outside the focal plane. An additional advantage of this system is that most optical components (e.g., laser, camera, mirrors) can be placed remotely on a vibration isolation table using commodity components. Therefore, concessions in performance do not need to be made to minimize weight, as only the fiber lens array is attached to the head of the animal, and optics can be readily re-configured for alternative imaging methods, such as dual-spectral reflectance imaging (see below).

**Figure 1.**
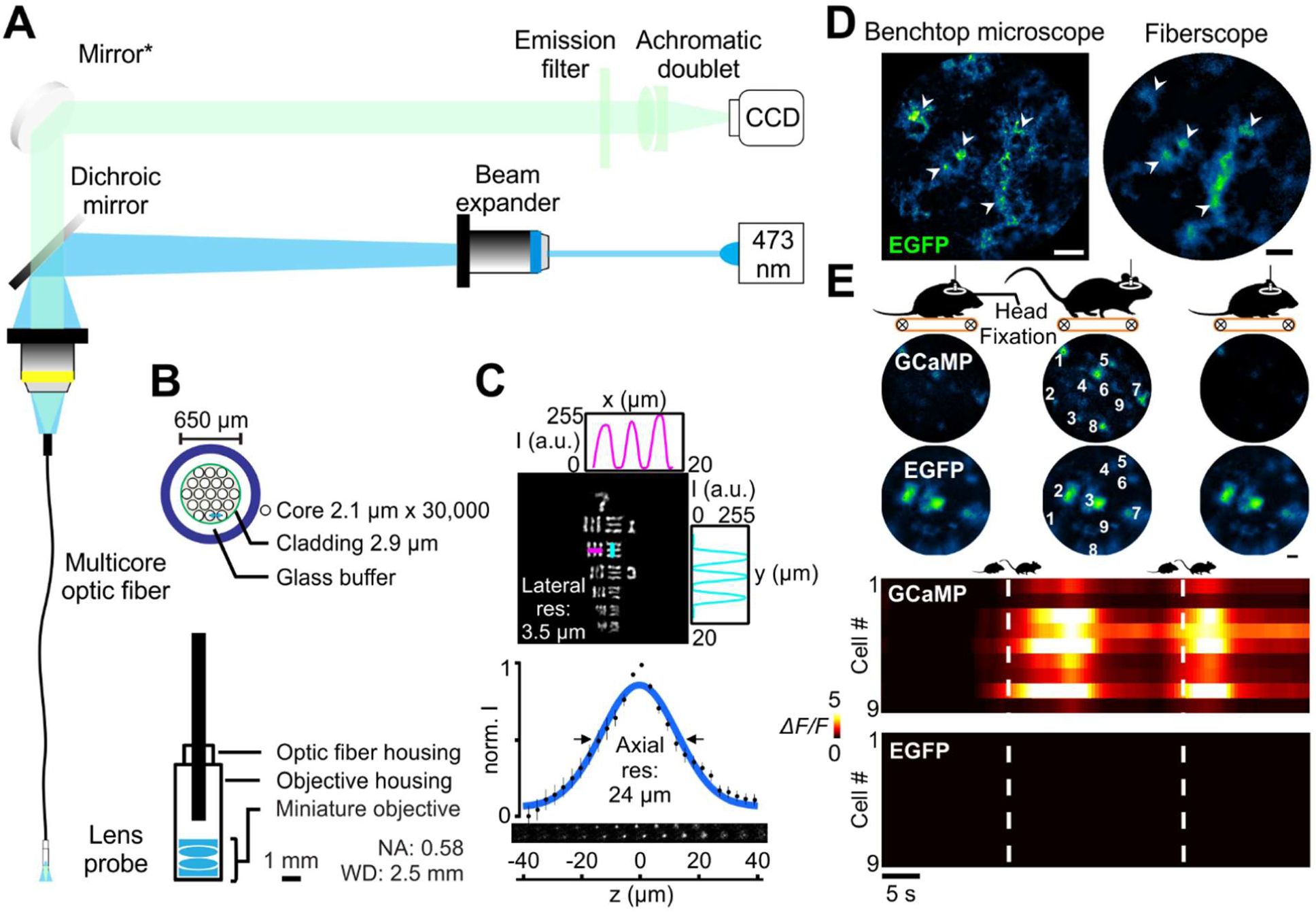
An optimized fiberscope for imaging astrocyte calcium dynamics *in vivo*. (A) Optical layout of the fiberscope imaging system. (B) Cross-section of multicore optical fiber bundle (top) and miniature objective configuration (bottom). (C) Resolution of the multicore fiber-miniature objective: 3.5 µm lateral resolution as in group 7 element 2 of the USAF target (top) and 24 µm axial resolution determined by the point-spread function of imaged microbead (bottom). (D) Comparison between a standard confocal microscope and the fiberscope imaging system using fixed tissue immuno-stained for EGFP. The white arrow heads denote individual astrocytes. Scale bar is 50 µm. (E) *In vivo* imaging using the fiberscope in head-fixed mice expressing either GCaMP or EGFP, and undergoing forced locomotion. Note that locomotion-induced increases in fluorescence were observed only in mice expressing GCaMP, indicating that these changes were not due to brain motion or other artifacts. Scale bar is 50 µm.

The imaging performance of this custom fiberscope was first benchmarked against a commercial confocal microscope. Images of cortical astrocytes expressing the genetically encoded calcium indicator GCaMP3 (*GLAST-CreER; Rosa26-lsl-GCaMP3* mice; astro-GC3 mice) (Paukert *et al*., 2014) in fixed brain slices were collected using a conventional confocal microscope and the fiberscope. This comparison revealed that the multicore system can resolve individual astrocytes and their processes (white arrowheads, Figure 1D), providing subcellular resolution similar to the benchtop system. To assess the performance of this system when imaging astrocytes *in vivo*, a micromanipulator was used to position the tip of the fiberscope over a cranial window implanted above the primary visual cortex (V1) of head-immobilized mice that expressed either GCaMP3 (astro-GC3) or EGFP (*GLAST-CreER; RCE:loxP* mice; astro-EGFP mice) selectively in astrocytes. Astro-EGFP mice served as a control for artifacts arising from movement or microenvironmental fluctuations, such as pH. In astro-EGFP mice the fiberscope was able to resolve single EGFP-expressing cells in V1, and in astro-GC3 mice it was able to detect GCaMP3 fluorescence within astrocytes in the imaging field (Figure 1E). Mice were induced to walk on a motorized treadmill (scheme, Figure 1E), a condition known to elicit a global rise in calcium within cortical astrocytes due to the local release of norepinephrine (NE) that accompanies enhanced arousal (Ding et al., 2013; Paukert *et al*., 2014; Reitman *et al*., 2023; Srinivasan *et al*., 2015; Streich et al., 2021). Astrocytes throughout the imaging field in astro-GC3 mice exhibited transient increases in fluorescence (Figure 1E) time-locked to the onset of ambulation, consistent with NE inducing global release of calcium from intracellular stores. Under comparable imaging conditions, no appreciable fluorescence changes were observed when astro-EGFP mice were induced to walk, indicating that the fluorescence changes in astro-GC3 mice arose though changes in intracellular calcium rather than motion-associated artifacts. Together, these results show that this lensed multicore fiber imaging system has sufficient sensitivity and resolution to monitor calcium levels within individual astrocytes *in vivo*.

### Imaging individual astrocyte activity in freely moving mice

To allow imaging of cortical astrocyte activity in freely moving mice, we designed a rigid, adjustable metal collet to secure the lensed-fiberscope to the mouse skull (Figure 2A). This mount allowed fine positioning of the fiber relative to the cranial window in both x-y and z planes, and facilitated longitudinal analysis of astrocyte activity over multiple days, as the fiberscope could be removed and readily repositioned to image the same focal plane. Astrocyte activity was monitored simultaneously with animal behavior by synchronizing the output of the fiberscope CCD with the output of two orthogonally-oriented near infrared (NIR) cameras placed outside the arena (Figure 2B). Animals that were allowed to roam freely in the arena with the fiber tethered exhibited similar behaviors as those that were untethered, in terms of their preferred location within the cage, the total distance moved (untethered: 0.068 ± 0.009 km, n = 7; tethered: 0.079 ± 0.008 km, n = 7; Student’s *t*-test, t (12) = 0.88, *p* = 0.40), and the speed of their movement (untethered: 5.7 ± 0.27 m/s, n = 7; tethered: 6.7 ± 0.37 m/s, n = 7; 2-sample *t*-test, t (12) = 2.1, *p* = 0.06) (Figure 2C).

**Figure 2.**
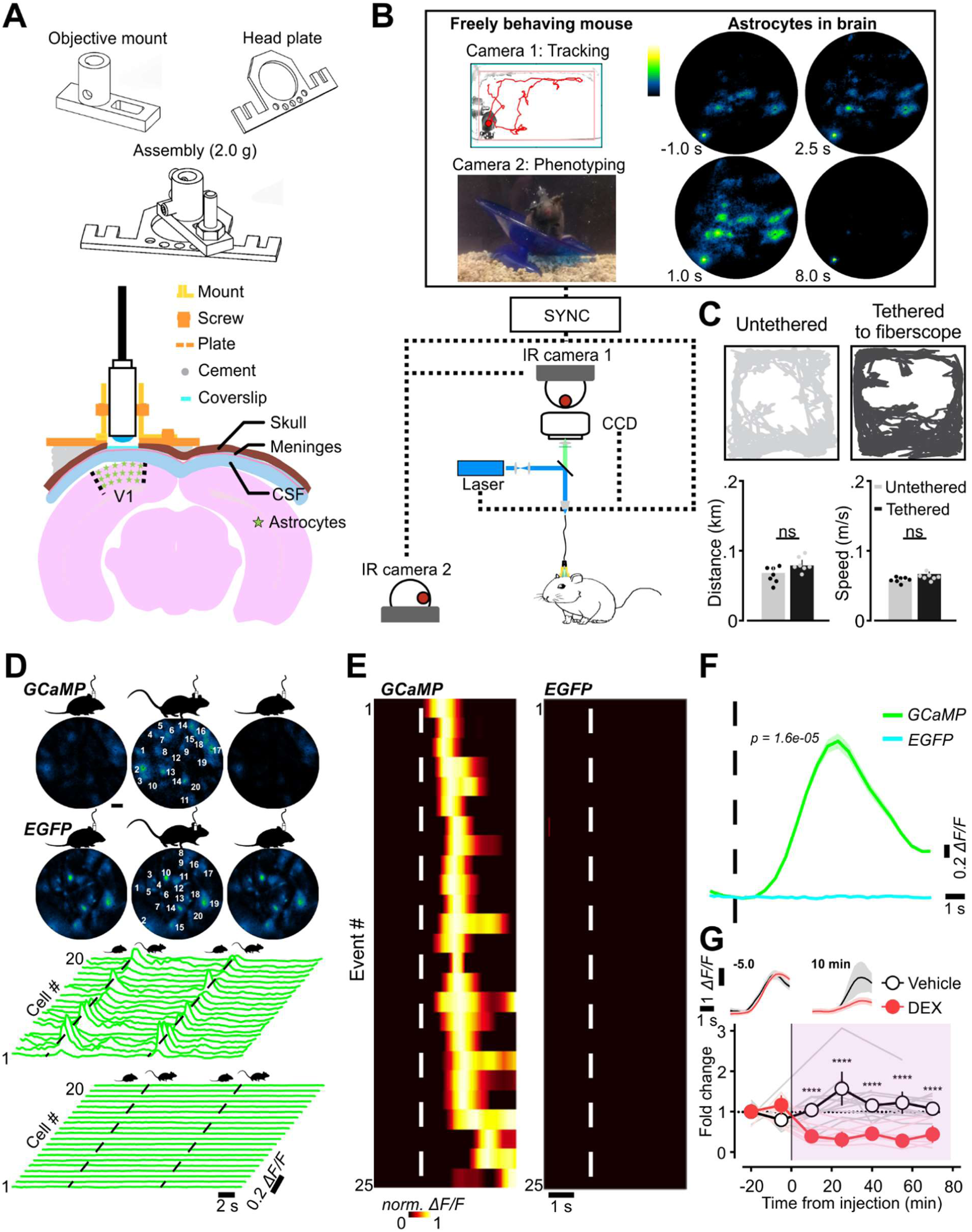
Imaging of astrocyte calcium in freely behaving mice. (A) Diagram of fiberscope-mouse brain coupling. Top: three-dimensional render of the mount for securing the fiberscope to the mouse skull. Bottom: minimally invasive optics-animal coupling for imaging of cortical astrocytes through cranial windows. (B) Outline of the configuration used to simultaneously monitor astrocyte calcium signals and mouse behavior. Output of the brain imaging CCD camera were synched with the two behavior-recording IR cameras. (C) Plots of mouse behavior over one hour showing that tethering to the optical fiber does not substantially change their behavior or the total distance they travel (n = 7 for each group). (D) Imaging fluorescence transients of 20 cells in a freely moving GCaMP and an EGFP animal. Corresponding traces quantified increases in GCaMP fluorescence (calcium increases) in astrocytes within the visual cortex during voluntary locomotion, in contrast to those of the EGFP animals where no significant changes were found. Scale bar is 50 µm. (E) Rank-ordered population (average of 20 cells) fluorescence intensity plot for 25 voluntary locomotion events for mice expressing GCaMP or EGFP. (F) Summary data for locomotion-induced fluorescence changes in astrocytes in mice expressing GCaMP or EGFP. (G) Plot of fold-increase in astrocyte fluorescence (calcium) during voluntary locomotion in mice injected with vehicle (1.14 ± 0.19, n = 6) vs. dexmedetomidine (0.33 ± 0.06, n = 9). Global locomotion-induced calcium transients are triggered by norepinephrine release, as they are strongly attenuated by dexmedetomidine, which inhibits the release of NE.

To determine if this configuration was stable enough to allow astrocyte imaging in freely moving mice, we monitored fluorescence changes in both astro-EGFP and astro-GC3 mice. When animals voluntarily transitioned from a quiescent to active state (Q-A transition), astrocytes in astro-GC3 mice exhibited consistent increases in fluorescence (Figure 2D), which occurred within ∼1 s after the onset of movement (dashed black line, Figure 2D), as shown by raster plots of across-cell fluorescence changes aligned to movement onset (dashed white line, Figure 2E), both for individual trials and when averaged across trials and animals (2.59 ± 0.59 Δ*F/F* at 90% rise; n = 6) (Figure 2F). In contrast, when similar experiments were performed in astro-EGFP mice, Q-A transitions did not elicit changes in astrocyte fluorescence (0.03 ± 0.01 Δ*F/F* at the time of 90% rise in astro-GC3 mice; n = 5) (Figure 2E, F). Astrocyte GCaMP activity associated with Q-A transitions was strongly suppressed when animals were administered the α2-adrenoceptor agonist dexmedetomidine (DEX, 3 µg/kg intraperitoneal), which binds to presynaptic receptors to inhibit release of NE from noradrenergic nerve terminals (Gelegen et al., 2014) (difference in fold change at 10 min: 0.81 ± 0.17; 95% CI, [0.3, 1.3]; Vehicle: n = 6, DEX: n = 9; two-way repeated measurement ANOVA, F (1, 90) = 111, *p* < 0.0001) (Figure 2G), indicating that astrocyte calcium levels are consistently elevated by NE during spontaneous arousal.

Neuronal and glial cell activity can cause local changes in blood volume, blood oxygenation, and changes in the optical properties of the brain (Devor et al., 2008; MacVicar, 1997; Rector et al., 2005), which alter the transmission of light and influence the detection of fluorescence signals (Kozberg et al., 2016; Ma et al., 2016). To assess the degree to which responses recorded from astro-GC3 mice are influenced by changes in neurovascular-associated absorbance, during *in vivo* imaging we measured the reflectance signal at 473 nm and 523 nm, corresponding to the peak excitation/emission wavelengths used for GCaMP imaging (Figure S1A). Measured GCaMP fluorescence signals were then corrected to compensate for these hemodynamic changes (Figure S1B) (Kozberg *et al*., 2016; Ma *et al*., 2016). Although Q-A transitions were associated with small reflectance changes at both wavelengths (Ex 473 nm, Em 523 nm) (Figure S1C), consistent with findings in head-restrained mice (Carandini et al., 2015), these changes in light absorbance had a minimal effect on the amplitude and time course of GCaMP fluorescence signals (Figure S1C). Hemodynamic-dependent cross-talk in GCaMP fluorescence signals depends on the path length travelled by photons. As most GCaMP fluorescence in this prep arises from astrocytes in the upper cortical layers, photons travel a shorter path to the fiber end, and are therefore contaminated exponentially less by hemodynamics than those from deeper layers (Hillman et al., 2011). Thus, GCaMP fluorescence changes in subsequent experiments were not adjusted to account for these hemodynamic reflectance changes, unless otherwise noted. Together, these studies indicate that the lensed multicore fiberscope is capable of resolving fluorescence signals from individual astrocytes during animal movement and is sufficiently sensitive to detect increases in GCaMP3 fluorescence initiated by physiological changes in cytosolic calcium.

### Longitudinal imaging reveals changes in astrocyte activation during the circadian cycle

Astrocytes have been shown to influence sleep homeostasis (Clasadonte et al., 2017; Ingiosi *et al*., 2020). In particular, sleep pressure, a phenomenon in which the motivation for sleep increases progressively during wakeful periods, is thought to be encoded by astrocyte calcium activity (Blum et al., 2021; Ingiosi and Frank, 2022). This accumulation of sleep need leads to release of astrocytic sleep-promoting substances such as adenosine (Choi et al., 2022). To understand how astrocytes are engaged to control this behavior, there is a pressing need for longitudinal, *in vivo* monitoring of astrocyte activity with cellular resolution during endogenous sleep-wake transitions. Therefore, we imaged freely moving mice during different phases of the circadian cycle (Figure 3A). To minimize photobleaching, the field-of-view was illuminated for 1 s every 60 s (Figure 3A). Sleep was inferred when the animal was immobile for >40 s, which has previously determined to have 95-99% correlation with simultaneous electroencephalography/electromyography-defined sleep (Pack et al., 2007; Fisher et al., 2012). Mice carrying the fiberscope exhibited gradual changes in movement over 24 hrs, evident when individual activity profiles were aligned to the most active period (Figure 3B), suggesting that this imaging approach allowed preservation of intrinsic circadian behavior. We then monitored individual astrocyte calcium levels over 24 hrs to define their spatiotemporal activity across the circadian cycle (Figure 3C). Within individual animals, the frequency of global astrocyte events (events that occurred in astrocytes throughout the imaging field) tracked with the phase of animal activity (Figure 3D), perhaps reflecting overall levels of neuromodulatory output. As expected, astrocyte calcium transients occurred more frequently during the active phase (0.21 ± 0.03, 95% CI, [13.7, 28.1], n = 7), than the inactive phase (0.07 ± 0.02, 95% CI, [2.89, 11.0], n = 7) (Wilcoxon rank-sum test, U = 47.0, *p* < 0.001) (Figure 3E). Similarly, when astrocytes were active, it was more likely that the animals were also active (Figure 3F; actual: 31.7 ± 0.1, 95% CI, [25.6, 38.7], n = 1000 *vs.* random: 24.0 ± 0.1, 95% CI, [19.9, 28.7], n = 1000) (2 sample K-S test = 0.8480, *p* < 0.001). These findings show that astrocyte calcium event frequency varies predictably with animal activity levels according to the circadian cycle, providing a means for astrocytes to integrate information about cumulative sleep and wake periods.

**Figure 3.**
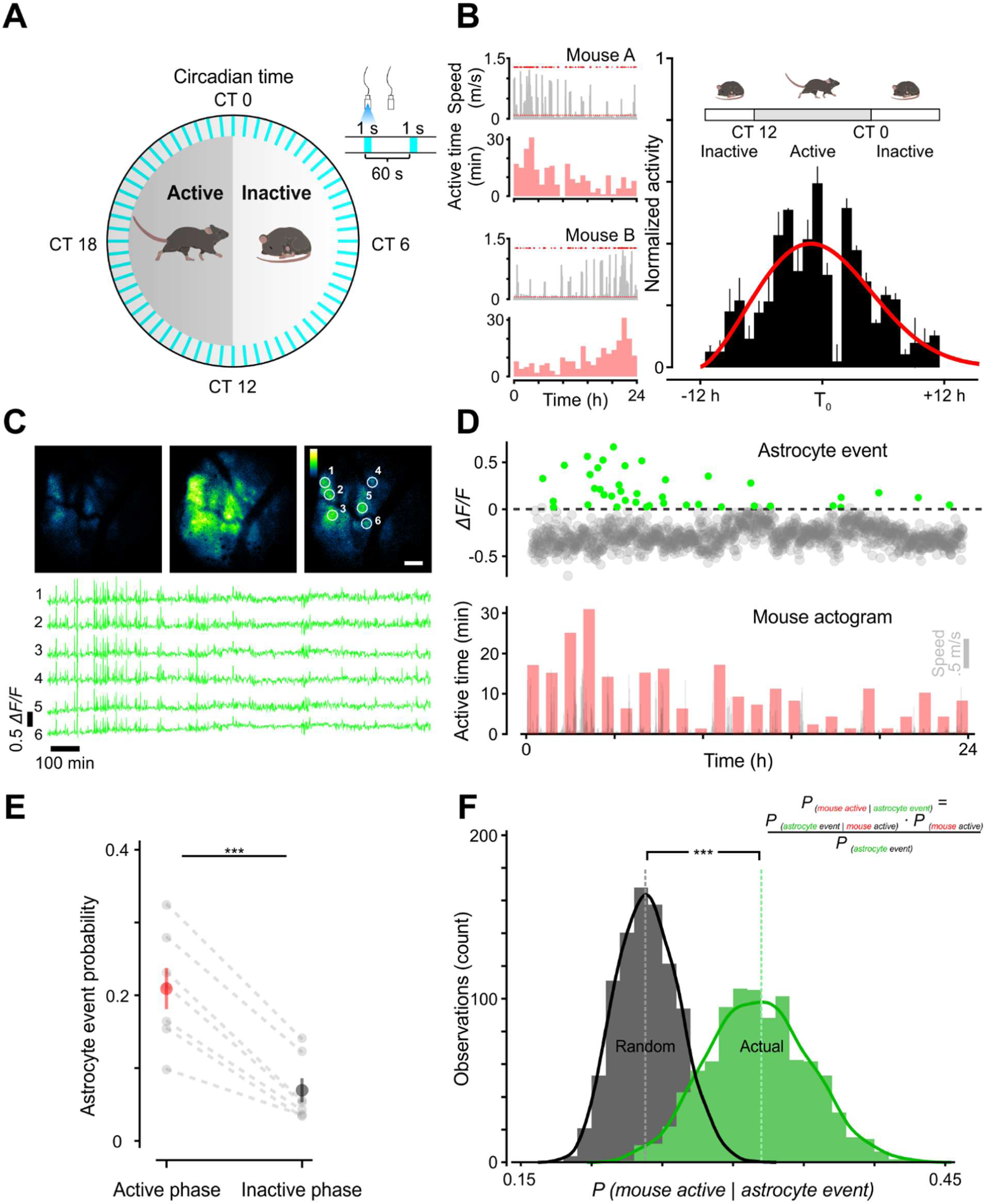
Freely moving imaging across 24 hrs reveals diurnal variation in astrocyte activity. (A) Scheme of astrocyte monitoring in a behaving animal across 24 hrs. (B) Left: two examples of activity level variations following the intrinsic rhythms of individual mice over 24 hrs. Right: phase-aligned hourly activity across animals (n = 6) showed robust preservation of the circadian cycle (red: probability distribution fitted curve). (C) Top: representative images from a 24-hour session showing astrocytes with baseline, maximal, and intermediate activity, respectively. Scale bar is 50 µm. Bottom: corresponding Δ*F/F* traces of the six cells highlighted in the image above. (D) Astrocyte calcium transients (green: astrocyte activation) appeared to follow the diurnal variation of mouse activity (bottom). (E) Astrocytes show more transients during the active phase of the diurnal cycle (red: 0.21 ± 0.03, n = 7) than during the inactive phase (black: 0.07 ± 0.02, n = 7) across all seven animals. (F) Astrocyte transients are indicative of animal activeness. Black denotes the Bayesian estimated probability if activity of the mouse were a random process; green represents the calculated probability from actual data (bootstrapped 1,000 times).

### Astrocytes are active during exploration rather than maintenance behaviors

A major advantage of imaging in freely moving animals is the ability to define which of a broad spectrum of behaviors are associated with astrocyte activity. To assess which behaviors were temporally correlated with changes in astrocyte calcium signaling, we monitored astrocyte fluorescence in the visual cortex of astro-GC3 mice as animals engaged in a variety of normal behaviors in addition to the Q-A transitions, such as grooming, nesting, exploring, drinking, and feeding (Figure 4A, B). To detect and annotate these behaviors in NIR video sequences, we used a machine learning-based detection program (Jhuang et al., 2010; Kabra et al., 2013) that was 94 ± 3.0 % accurate in categorizing these behaviors, as independently assessed when blinded to the computer-aided categorization (see Methods) (Figure 4A). Although transient increases in astrocyte calcium levels were occasionally observed during diverse behaviors, Q-A transitions and exploratory behaviors such as rearing were most prominently correlated with enhanced astrocyte activity (Figure 4B, C). These studies are consistent with the conclusion that astrocyte calcium transients occur most prominently during periods of enhanced arousal, though not exclusively during locomotion.

**Figure 4.**
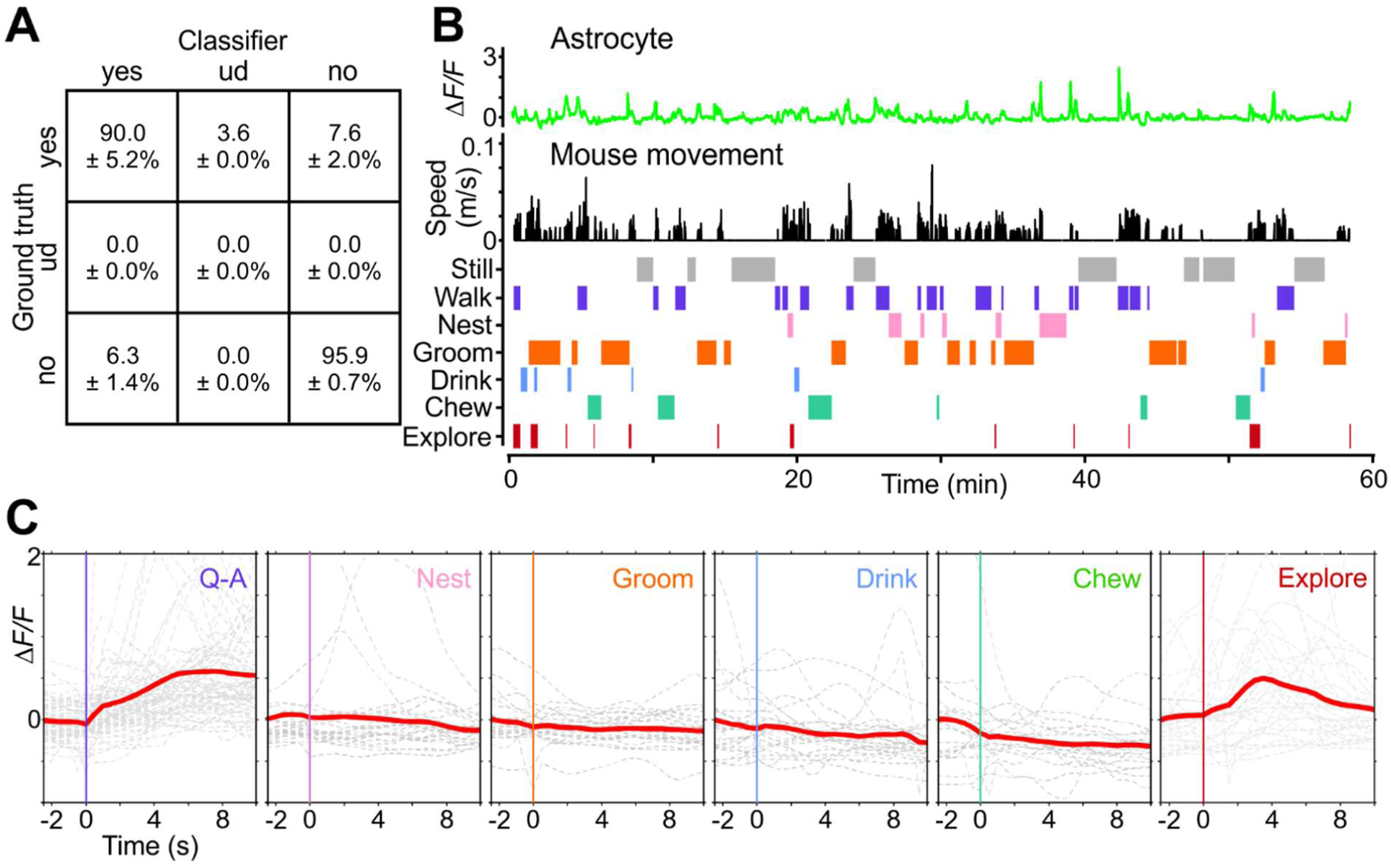
Astrocytes exhibit distinct calcium responses during different behaviors. (A) Human-provided ground-truth labels over 50,000 frames yielded 93.9 ± 3.0% accuracy for the machine learning classifier in categorizing behaviors (ud: undetermined). (B) Longitudinal monitoring of astrocyte calcium changes (top) and mouse movements (middle) during spontaneous behaviors (bottom). (C) Plots of astrocyte calcium transients aligned to the onset of behavioral events. Gray lines are individual events and red lines are the average of all events. Note that the most consistent activation occurred at Q-A transitions and during exploration, such as rearing.

### Diversive exploration in novel environments activates astrocyte networks

Animals often exhibit characteristic exploratory behavior when encountering a new environment (Berlyne, 1950; Glickman and Sroges, 1966; Welker, 1957). For example, mice respond to new environments by rearing on their hind legs to gather information about their surroundings (Lever et al., 2002), a diversive explorative behavior associated with spatial/contextual learning (Kemp et al., 2004; Xu et al., 1998). We therefore investigated whether calcium signaling was induced in astrocytes when mice explored novel areas (Lever et al., 2006). To quantify astrocyte activity explicitly during this behavioral context, we identified rearing events from movies as mice explored the arena by continuously computing body length (see Methods), which becomes highly extended in mice during rearing. This algorithm reliably distinguished rearing from other movements in the arena (Figure 5A, C), allowing GCaMP fluorescence signals in astro-GC3 mice to be aligned to each rearing event (Figure 5B, C). Analysis of these events revealed that rearing was consistently associated with increases in astrocyte calcium levels, visible as transient increases both in the GCaMP fluorescence of individual astrocytes in an imaging field (Figure 5B), and in the average cell fluorescence change across animals (Figure 5D); in contrast, animals that were continuously active, but not rearing, did not exhibit consistent changes in astrocyte calcium (Figure 5C, D). Correction of these fluorescence measurements for hemodynamic changes revealed that rearing events were consistently associated with increases in astrocyte calcium (Figure S1C); however, the amplitude of the GCaMP fluorescence rise was not obviously correlated with either the duration of the rearing event (Δt _rear_) or the magnitude of body extension (Δh _rear_) (Figure 5E).

**Figure 5.**
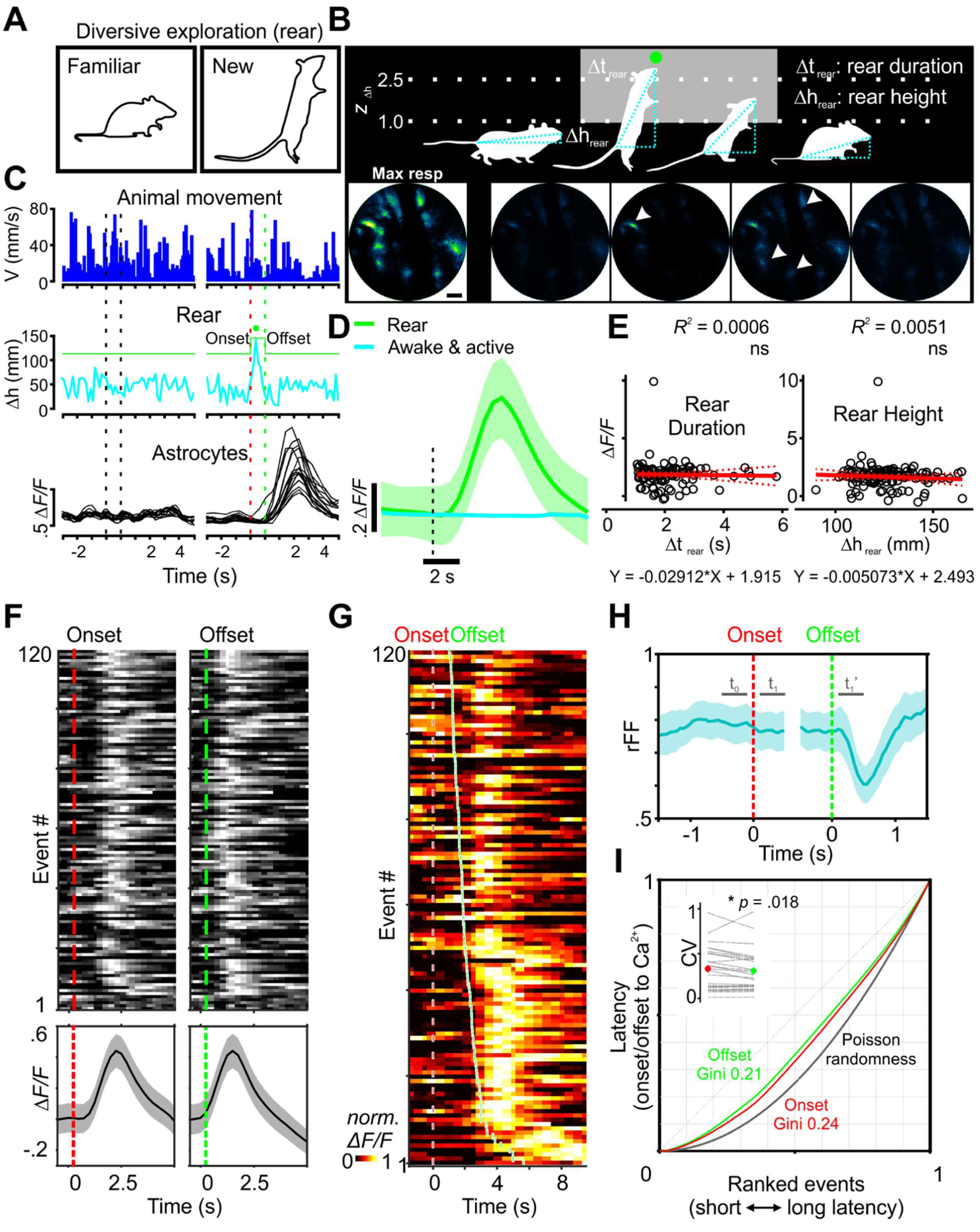
Diversive exploration (rearing) in environmental novelty engages astrocyte networks. (A) Diagram of encouraged rearing, a selective form of diversive exploration, in novel spatial environments. (B) Top: quantitative analysis for capturing and parameterizing each rearing event. Bottom: representative GCaMP fluorescence images in astro-GC3 mice during rearing. Scale bar is 50 µm. (C) Traces of cortical astrocyte activities (black) when mice moved continuously (blue) and reared (cyan). (D) Calcium transients plotted across animals, comparing rearing to moving behaviors. (E) Transient magnitude was not correlated with duration of the rearing event (Δt _rear_) or the extent of body extension (Δh _rear_). (F) GCaMP fluorescence was aligned (heat map) and averaged (bottom trace) across events to the onset and offset of the behavior. (G) Onset-aligned and offset-ranked raster plot demonstrated a concomitant increase in the delay of the peak in astrocyte calcium when rearing events lasted longer. (H) During rearing events, in comparison with baseline (t_0_, denoting the final 900 ms to 100 ms before the event), the rFF reduced as little as 5.7% after rearing onset (t_1_, from 100 ms to 900 ms after), but declined by 21.6% after the end of offset (t_1_’, from 100 ms to 900 ms after), indicating a closer association between astrocyte calcium and rearing offset. (I) Lorenz plots for the latency of the peak astrocyte calcium transients and the onset and offset times: Gini was 0.24 for onset latency distribution and 0.21 for offset. Inset: coefficient of variation (CV) of latencies to the astrocyte response across animals was higher for the onset (red: 0.32 ± 0.05, n = 25) than for the offset (green: 0.30 ± 0.05, n = 25).

To understand which aspects of this complex behavior were most closely linked to astrocyte calcium changes, the peak increase in GCaMP fluorescence was aligned to the onset and offset of the behavior and rank-ordered relative to delay time (Figure 5F). Because the peak fluorescence change occurred with a latency of 2.45 ± 0.12 s after onset (n = 120), and rearing events followed a relatively consistent time course lasting 1.53 ± 0.07 s (n = 120), the maximal astrocyte calcium response occurred well after onset of the behavior (Figure 5F, G). When rearing events lasted longer, there was a concomitant increase in the delay to the peak in astrocyte calcium, visible when onset and offset time were included on the same rank-order raster plot (Figure 5G), suggesting that astrocyte activity is more closely linked to the cessation rather than the onset of this behavioral sequence. The consistency in the timing between this behavioral sequence and peak astrocyte calcium was assessed further by calculating the rFF (Fano factor, the ratio of the variance to the mean of the calcium transients among defined intervals), which is often used to describe the variance of action potentials in spike trains during *in vivo* recordings (Nandy et al., 2017; Nawrot, 2010), in which lower rFF values indicate higher deviation from a random process (Lee et al., 2017; Montijn et al., 2014). During rearing events, the rFF deviated 5.7% from baseline upon rearing onset (Figure 5H, t_1_, from 100 ms to 900 ms after baseline), but declined as much as 21.6% at the offset (Figure 5H, t_1_’, from 100 ms to 900 ms after), indicating a closer association between astrocyte calcium and rearing offset. To better quantify this relationship and address the possibility of random association, we analyzed the distribution of latency from the peak astrocyte transient to the time of onset and offset using Lorenz plots with Gini coefficient (Yu et al., 2019), providing a measure of how these intervals vary from an equal distribution. The Gini coefficient was 0.24 for the onset latency distribution and 0.21 for the offset distribution (Figure 4I). The difference in Gini coefficients indicates that the associations are not random and that there is a closer relationship between the peak and event offset than event onset. Moreover, the coefficient of variation of latencies to the peak astrocyte response was slightly higher for the onset (0.32 ± 0.05, 95% CI, [0.23, 0.42], n = 25) than for the offset (0.30 ± 0.05, 95% CI, [0.20, 0.39], n = 25) (Wilcoxon signed-rank test, *p* = 0.02) (Figure 5I, inset). Thus, as compared to enforced locomotion and spontaneous Q-A transitions that elicited rapid elevation of calcium, in this exploratory behavior, astrocyte networks were consistently activated following the end of the behavioral sequence, indicating that astrocytes can be activated during discrete periods of complex behaviors.

### Astrocytes become desynchronized and adapt rapidly during inspective exploration

We further assessed astrocyte activity when animals inspected novel objects (Figure 6A, scheme). Mice that demonstrated no more than 55% object bias in a familiarization session proceeded to the test session (*p* = 0.48 for percentage of time spent exploring two familiar objects, F or F’) (Figure 6A). During test sessions, mice were returned to the arena where one of the familiar objects had been replaced with a new object. All animals in test sessions spent more time exploring the new object (percentage of time spent exploring the familiar object, F: 24% ± 3.5%, 95% CI, [16%, 32%] or the new object, N: 76% ± 3.5%, 95% CI, [68%, 84%], n = 10 animals) (paired *t*-test, t (9) = 7.5, *p* < 0.0001) (Figure 6A), indicating that they recognized novelty in the current setting (discriminability index d’: 0.52 ± 0.07, 95% CI, [0.37, 0.68]) (1-sample *t*-test, t (9) = 7.5, *p* < 0.0001). We then monitored calcium fluctuations in astrocytes within the visual cortex while animals explored either novel or familiar objects (Figure 6B). The time of onset of directed exploration was defined as the moment the mouse came within 2.5 cm and oriented their head towards the object (Figure 6B, green dashed lines). Inspective exploration of the novel object triggered small, rapidly rising and decaying calcium transients in subsets of astrocytes that often began before onset of interaction (Figure 6B, *Novel*). In contrast, exploration directed toward a familiar object consistently failed to elicit detectable calcium increases in astrocytes (Figure 6B, *Familiar*).

**Figure 6.**
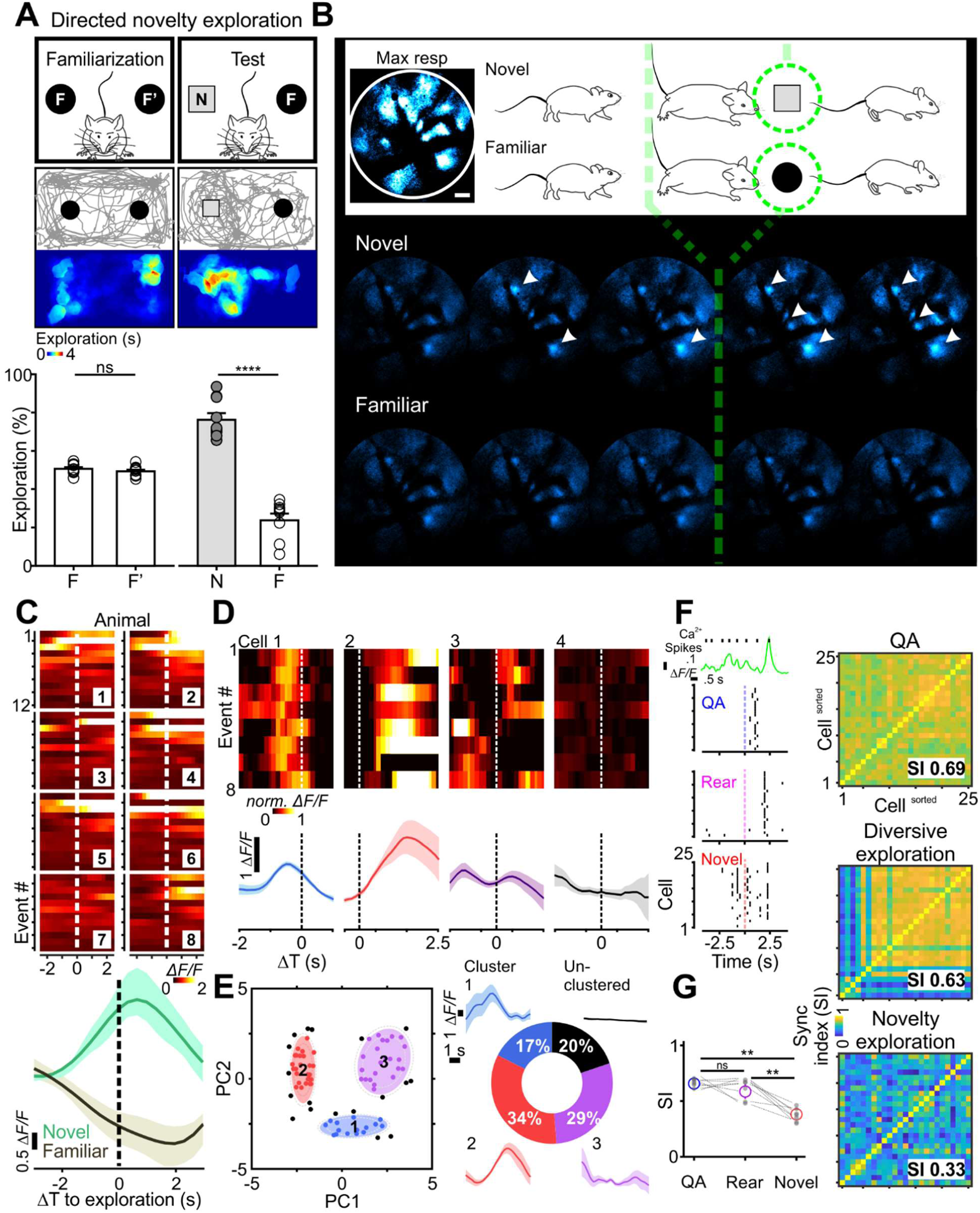
Astrocyte networks desynchronized and adapted rapidly during inspective exploration. (A) Scheme of novel object recognition paradigm (top) where a mouse preferentially traveled (trace) and spent time (color map) around the novel object. Bottom bar chart: establishment of paradigm across 10 animals, shown by consistent exploration preference toward the novel object. (B) Representative images (scale bar is 50 µm) of cortical astrocytes while animals explored the novel *vs.* the familiar object, with the onset of directed exploration indicated by green dashed lines. (C) In 8 representative animals, population transients (across-cell average) were time-locked to the directed exploration toward the novel object, but not toward familiarity. Of note, levels of astrocyte engagement declined rapidly with repetitive interactions, indicating fast adaptation from event 1 to 12. (D) Top: heat map with Δ*F/F* normalized to the max (norm. Δ*F/F*, ranging from 0 to 1) during each event suggested that timing of calcium transients within individual cells was remarkably consistent in relation to the behavioral sequence. Bottom: across-event calcium average of the individual cells. (E) Left: Gaussian mixture model with expectation maximization sorted principal component (PC1 and PC2) into three clusters (80.2% of 86 astrocytes) with the color edge at 95% likelihood (outer broken grey line: 99% likelihood). Right: within-cluster calcium averages suggested astrocytes of each cluster shared distinct temporal features. (F) Left: representative network of 25 cells during a Q-A transition, a diversive rearing exploration, and inspective novelty exploration (detected calcium transients plotted as black vertical raster). Right: synchronization indices to quantify the network difference visualized in the raster plots. (G) Across 8 animals, inspective explorations consistently desynchronized cortical astrocytes, distinct from the homogeneity observed when mice entered a state of enhanced arousal during Q-A transitions and rearing.

Quantifying full-field population dynamics across events for different animals (Figure 6C, color maps) revealed that astrocyte calcium signaling increased consistently during initial interactions with a novel object (0.29 ± 0.08 Δ*F/F* at 90% rise; 95% CI, [0.04, 0.56], n = 8) (Figure 6C), but not during interactions with a familiar object (–0.24 ± 0.06 Δ*F/F*; 95% CI, [–0.65, 0.14], n = 8) (Figure 6C), with the two conditions being significantly different (two-way repeated measures ANOVA, F (1, 7) = 5.5, *p* = 0.03). In accordance with behavioral adaptation, astrocytes exhibited a consistent decrease in responsiveness with successive interactions with a novel object (compare responses during the initial events to the later in the color map of Figure 6C); comparing the mean response (90% rise) of the last three events (–2.6 ± 0.7 Δ*F/F*, 95% CI, [–4.2, -1.0], n = 8) to that of the first three events (14.1 ± 5.9 Δ*F/F*, 95% CI, [0.2, 28.1], n = 8), revealed that the response of astrocytes decreased by 156 ± 20% (95% CI, [109%, 202%], n = 8). These findings indicate that cortical astrocyte activity is updated rapidly and adapts to changing behavioral contexts.

Unlike the coordinated astrocyte activity observed during Q-A transitions and rearing, astrocyte responses appeared much less correlated during directed novelty exploration (Figure 6B). To determine if individual astrocytes exhibited consistent responses during this behavior sequence, we aligned the GCaMP traces from individual cells (Figure 6D). This presentation revealed that despite different response profiles, within individual astrocytes the peak of calcium changes with respect to the time of object encounter seemed remarkably consistent across sessions. We attempted to cluster astrocytes according to the timing and magnitude of their peri-exploration calcium transients over eight consecutive trials. Applying principal component analysis to the data yielded a first component containing 76.6% of the variance, and a second containing 12.2% of the variance. A Gaussian mixture model was then fitted to the scores of these two components using an expectation-maximization algorithm (see Methods), which, by threshold of 95% likelihood, identified three main subpopulations of astrocytes (of 86 astrocytes analyzed) with shared response characteristics (Figure 6E, Cluster 1-3). The average calcium transients for astrocytes were plotted for each cluster, revealing that 80.2% of astrocytes maintained their distinct temporal features between trials (Figure 6E, traces). The temporal differences in their responses enabled astrocytes to, in aggregate, spread their activity over the course of the exploration event, with different groups aligned to discrete behavioral contexts (Figure 6C). Astrocytes that were not contained within the clusters (Figure 6E, black) typically exhibited responses that were inconsistent from trial to trial. Since clustering was based on calcium changes associated with peri-inspective exploration, these unclustered cells may serve as reserve to encode alternative behavioral contexts, as proposed for neurons.

To further define individual cellular activities in diverse contexts, we plotted calcium events from a representative network of 25 astrocytes (Figure 6F, detected calcium transients plotted as black vertical raster lines) during Q-A transitions, diversive rearing explorations, and directed novelty explorations (Figure 6F). This analysis revealed that directed explorations were associated with less synchronized astrocytic activity (Figure 6F, Novel), in contrast to the highly-synchronized calcium events observed when the mouse experienced a state of arousal (Figure 6F, QA, SI = 0.69), a phenomenon seen for all animals (Figure 6G, 0.67 ± 0.02, 95% CI, [0.64, 0.71], n = 8). Desynchronization emerged when the mouse explored new surroundings (Figure 6F, Rear, synchronization index (SI) = 0.63), and became most prominent when its attention was directed toward a target (Figure 6F, Novel, SI = 0.33). This astrocyte desynchronization in directed/inspective exploration was significant across animals (Figure 6G, 0.36 ± 0.03, 95% CI, [0.30, 0.43], n = 8) (Friedman test, F (9) = 12, *p* < 0.01). Although a form of exploration, rearing did not produce such marked desynchronization (Figure 6G, 0.63 ± 0.04, 95% CI, [0.53, 0.73], n = 8), perhaps because rearing is often associated with increased vigilance/arousal (Lever *et al*., 2006).

### Contextual encoding by cortical astrocytes is dependent on convergent neuromodulation

To explore the neural mechanisms responsible for the distinct response profiles of astrocytes in different behavioral contexts, we monitored changes in their responses after topical application of antagonists to the cortical surface (Figure 7A, scheme). In accordance with previous studies (Figure 2G, Paukert et al., 2014), the α1 adrenoceptor antagonist prazosin reduced the amplitude of astrocyte calcium transients induced during Q-A transitions (Δ*F/F _after-before_*: aCSF 0.00 ± 0.04, n = 10 *vs.* prazosin 0.25 ± 0.09, n = 8) (Kruskal-Wallis, H = 18.4, *p* = 0.001) (Figure 7A, Q-A); in some experiments, prazosin also slowed the rise of responses, but this did not reach significance (*p* = 0.06). Astrocyte responses during rearing events were similarly suppressed by prazosin (Δ*F/F _after-before_*: –0.19 ± 0.03); these events were also slower to reach the peak (Δt _from rear to 10% rise_: 0.45 ± 0.09 s) and shorter in duration (FWHM: –0.15 ± 0.45 s, n = 9) as compared to aCSF control (–0.08 ± 0.08 s, 0.06 ± 0.03 Δ*F/F*, –0.09 ± 0.35 s, n = 8) (one-way ANOVA for Δt _from rear to 10% rise_, F (4, 38) = 5.5, *p* = 0.01; Kruskal-Wallis for Δ*F/F _after-before_*, H = 12.7, *p* = 0.01; one-way ANOVA for FWHM, F (4, 38) = 3.4, *p* = 0.02) (Figure 7A, Rear). Together, these results indicate that α1 adrenoceptors play a critical role in engaging astrocyte networks in behaviorally relevant contexts associated with increases in arousal/vigilance.

**Figure 7.**
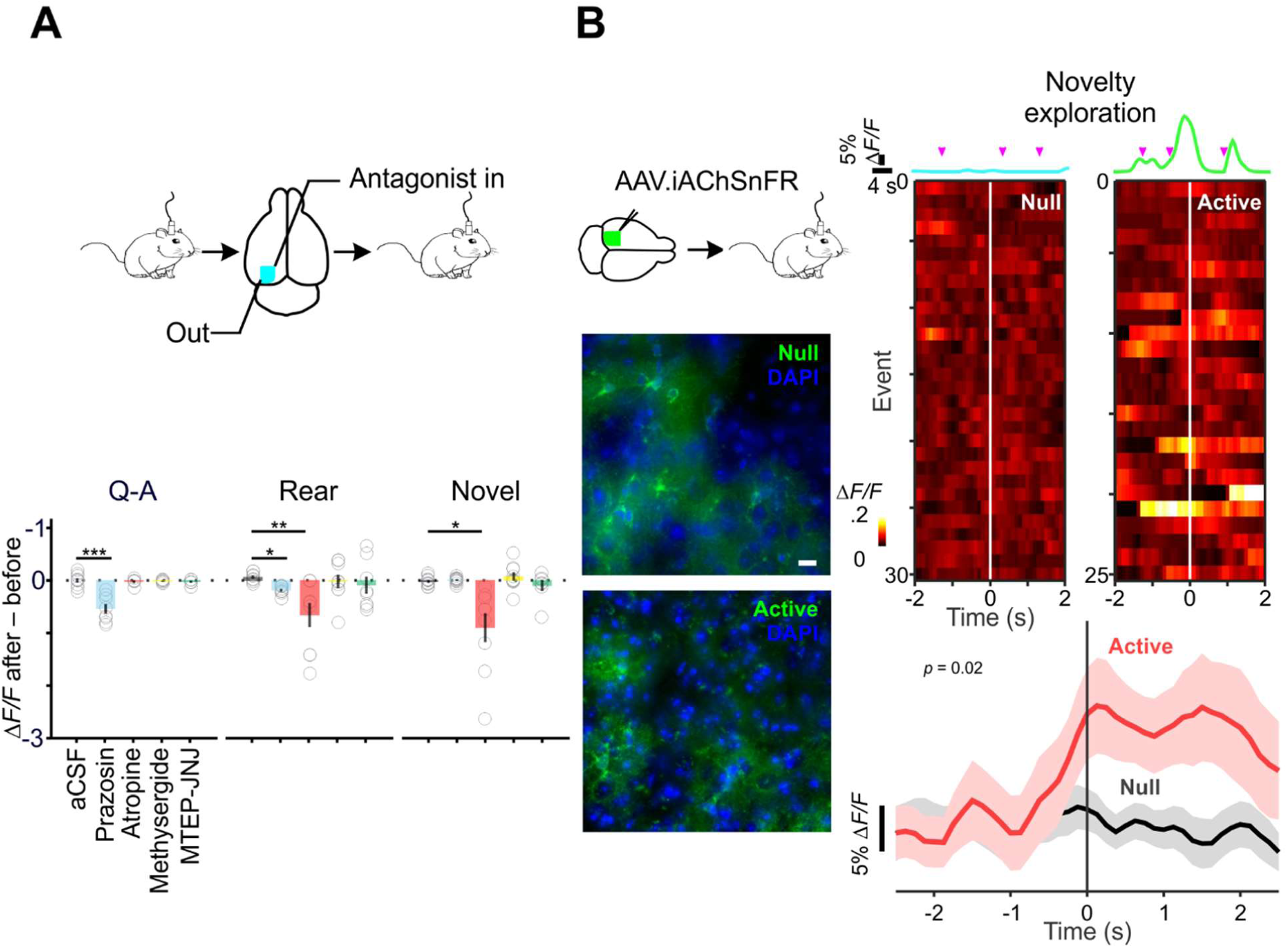
Divergent contextual encoding in cortical astrocytes depends on convergent neuromodulation. (A) Top: scheme of topical pharmacology in freely behaving animals. Bottom bar chart: prazosin altered all aspects of QA-relevant astrocyte response and diversive spatial exploration; atropine had a profound effect on amplitudes in contexts of exploration, be it diversive or inspective. (B) Left top: scheme of stereotactic introduction of *AAV.GFAP.iAChSnFR ^Active^* and *AAV.GFAP.iAChSnFR ^Null^* into the mouse cortex. Left bottom: immunofluorescent images of cortical astrocytes expressing *iAChSnFR ^Null^* (“Null”) and *iAChSnFR ^Active^* (“Active”); the preservation of fine ramification indicated no overt pathological changes in transduced astrocytes (scale bar is 50 µm). Right top: representative traces showing directed exploration reliably elicited fluorescence transients in the brains of iAChSnFR ^Active^ animals but not in those of the iAChSnFR ^Null^ ones. Right bottom: fluctuations aligned to each directed exploration (color raster) and summarized across animals (red: iAChSnFR ^Active^, n = 11 *vs.* black: iAChSnFR ^Null^, n = 12).

Previous studies have shown that electrical stimulation of nucleus basalis or direct application of ACh *in vivo* can elicit calcium increases in cortical astrocytes (Chen et al., 2012; Takata et al., 2011), and single-cell RNAseq analysis of astrocytes in the visual cortex indicate that they express muscarinic receptor M1 (*Chrm1*) (Bayraktar *et al*., 2020; Hrvatin et al., 2018), but the physiological contexts under which these receptors are activated is unknown. To test whether muscarinic receptors contribute to these astrocyte calcium changes, we locally applied the muscarinic antagonist atropine and measured their responses during different behavioral sequences. Atropine significantly reduced calcium transient amplitudes during both diversive exploration (–0.66 ± 0.23 Δ*F/F*, n = 9 *vs.* aCSF: 0.06 ± 0.03, n = 8) (Kruskal-Wallis, H = 12.7, *p* = 0.01) (Figure 7A, Rear) and inspective exploration (–0.90 ± 0.27 Δ*F/F*, n = 9 *vs.* aCSF: –0.02 ± 0.03, n = 11) (Kruskal-Wallis, H = 18.1, *p* = 0.001) (Figure 7A, Novel). In contrast, local inhibition of 5-HT_1/2_ (Methysergide) or mGluR_1/5_ (MTEP-JNJ) signaling did not significantly alter the characteristics of astrocyte responses. However, during inspective exploration, inhibiting mGluR_1/5_ enhanced the synchrony of astrocyte activation (MTEP-JNJ SI _after-before_: 0.06 ± 0.02), an effect opposite to that of α1 adrenergic antagonism (prazosin SI _after-before_: –0.10 ± 0.04) (one-way ANOVA, F (4, 42) = 2.9, *p* = 0.03). These findings suggest that cholinergic signaling in astrocytes is engaged during particular exploratory events, and that the coordination of astrocyte calcium transients can be altered by signaling through metabotropic glutamate receptors.

As an alternative method to explore whether astrocytes are exposed to ACh released from cholinergic terminals in specific behavioral contexts, we expressed a fluorescent ACh sensor selectively on the surface of astrocytes and monitored fluorescence changes during novelty exploration. The genetically encoded ACh sensor iAChSnFR (Borden et al., 2020) was expressed on cortical astrocytes by stereotactic delivery of *AAV.GFAP.iAChSnFR ^Active^* in V1 (see Methods); control mice were injected with the non-functional variant *iAChSnFR ^Null^* (Borden *et al*., 2020) using the same approach (Figure 7B, scheme). Viral delivery of these transgenes resulted in sparse, but specific, expression of the null and active sensors on astrocytes (Figure 7B, panels below). For mice engaged in novel object-directed exploration, each inspection event (Figure 7B, magenta arrow heads) elicited fluorescence changes in astrocytes expressing iAChSnFR ^Active^ but not iAChSnFR ^Null^ (Figure 7B, traces). These fluctuations were aligned to each directed exploration (Figure 6B, color raster) and summarized across animals (Figure 6B, red: iAChSnFR ^Active^, n = 11 *vs.* black: iAChSnFR ^Null^, n = 12). The significant change seen in iAChSnFR ^Active^ animals (Figure 7B, at 90% rise of iAChSnFR ^Active^, Δ*F/F* _Active – Null_ = 0.44 ± 0.24, 95% CI, [0.05, 0.58]) (mixed-effect model, F (1, 21) = 5.977, *p* = 0.02) indicates that ACh released in the cortex during novelty exploration (Acquas et al., 1996; Giovannini et al., 1998; Miranda et al., 2000) reaches astrocyte membranes at a level sufficient to bind to and elicit prolonged increases in fluorescence from this sensor.

### Direct modulation of astrocyte signaling by internal state

Astrocytes exhibited few calcium transients when engaged in maintenance behaviors, such as drinking, eating, grooming and nesting under *ad libitum* conditions (Figure 4C). However, these fundamental behaviors are also embedded with a motivational component, where the rewarding properties of resources can be magnified through deprivation (Cabanac, 1971; Toates, 1986). To examine the effects of internal homeostatic state on astrocyte responses to nutritional cues, we imaged the same mice across water- or food-restricted *vs.* naïve states (scheme, Figure 8A). Consistent with observations made during home cage monitoring (Figure 3C), astrocyte activation was not detected in satiated animals (water: Δ*F/F* –0.12 ± 0.05 in 24 mice, food chow: Δ*F/F* –0.19 ± 0.08 in 20 mice) (Figure 8A). Strikingly, in animals subjected to water deprivation (Δ*F/F* 0.47 ± 0.12 in 17 mice) or food deprivation (Δ*F/F* 0.46 ± 0.21 in 22 mice), astrocytes were robustly activated when mice were exposed to these previously neutral cues (Figure 8A). However, this increase was only significant in water-deprived conditions, due to high variability among animals (difference of Δ*F/F* in water-deprived *vs.* satiated animals: 0.21, [0.01, 0.41]; in food-deprived *vs.* satiated: 0.18, [–0.02, 0.39]) (water: Wilcoxon rank-sum test, U = 128, *p* = 0.04; food chow: Wilcoxon rank-sum test, U = 154, *p* = 0.10) (Figure 8C). Examination of the response profiles of astrocytes revealed that their initial response occurred prior to drinking (calcium peak time: –1.5 s) or ingesting food (–1.75 s). Thus, we hypothesized that these deprivation-induced changes were triggered primarily by cue detection rather than consumption. To test this hypothesis, mice were presented with inaccessible cues (scheme, Figure 8B) in which they could see or smell food, but not engage with it. Simply detecting the cues without direct interaction resulted in rapid activation of astrocytes (smell: Δ*F/F* 2.3 ± 0.7 in 22 mice; see: Δ*F/F* 1.6 ± 0.5 in 25 mice) (Figure 8B). Here, the difference between the food-deprived and the satiated were much more striking (Δ*F/F* _deprived - satiated_ in smelling inaccessible food: 0.80, [0.33, 1.41], Wilcoxon rank-sum test, U = 51, *p* < 0.0001; Δ*F/F* _deprived - satiated_ in seeing: 0.67, [0.41, 1.18], Wilcoxon rank-sum, U = 66, *p* < 0.0001) (Figure 8C). Of note, these responses were much larger in magnitude than when they were allowed to eat (Kruskal-Wallis, H = 16.3, *p* = 0.0002), suggesting that the inability to obtain the goal may unmask and/or provoke greater neuromodulatory release as a consequence of enhanced and sustained awareness. Together, these findings indicate that the engagement of cortical astrocytes is not fixed within different behavioral contexts, but instead can be adjusted based on changes in internal motivational state.

**Figure 8.**
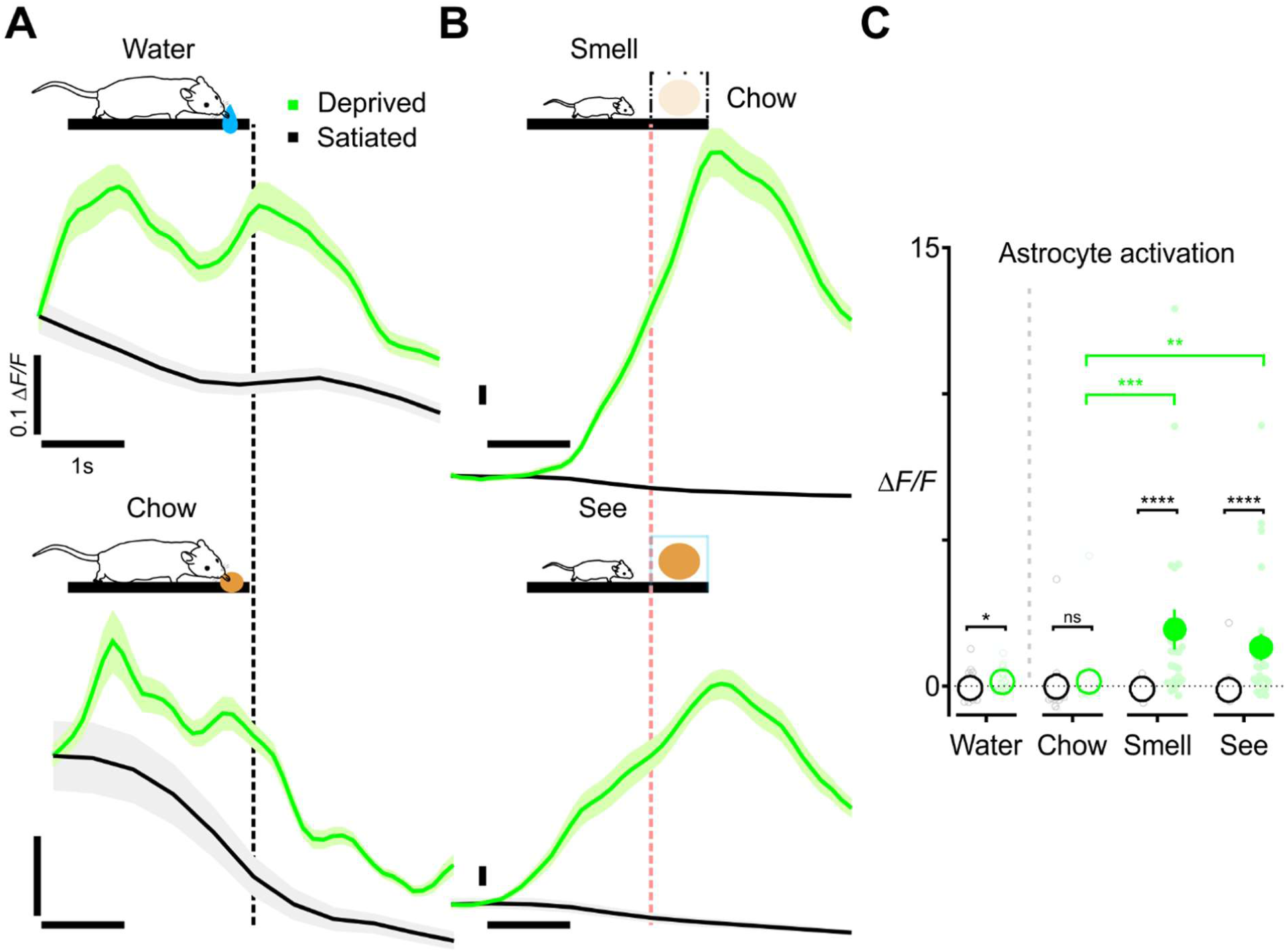
Internal state-driven astrocyte signaling of nutritional cues. (A) Imaging the same animal across water- (top) or food-deprived (bottom) *vs.* satiated states; astrocyte activation was not seen in metabolically-satiated animals (water: Δ*F/F* -0.12 ± 0.05, food: Δ*F/F* -0.19 ± 0.08). The population became sensitive to cues during behavioral states of thirst (Δ*F/F* 0.47 ± 0.12) or hunger (Δ*F/F* 0.46 ± 0.21). (B) Inaccessible cues augmented astrocyte activation (smell: Δ*F/F* 2.3 ± 0.7; see: Δ*F/F* 1.6 ± 0.5) compared to those of the accessible, a phenomenon that was not seen in satiated animals (satiated-smell: Δ*F/F* -0.02 ± 0.01; satiated-see: Δ*F/F* 0.01± 0.01). (C) Consistent astrocyte activation to cues in nutritionally-deprived animals across accessible and inaccessible cues (black asterisks) and augmented responses to inaccessible cues (green asterisks, smell *vs.* eat: *p* = 0.0007; see *vs.* eat: *p* = 0.003).

## Discussion

Astrocytes are now recognized as both a target of neuromodulatory influence and an effector that can induce broad changes in neural activity during different behavioral states (Chaturvedi *et al*., 2022; Doron *et al*., 2022; Mu *et al*., 2019; Srinivasan *et al*., 2015; Yu *et al*., 2020), but the relationship of their activity patterns to diverse naturalistic behaviors remains to be fully defined. The increasing recognition that cortical astrocytes vary in their molecular and physiological properties, between brain regions (Chai et al., 2017), cortical layers (Bayraktar *et al*., 2020; Miller *et al*., 2019), and within domains of particular signaling such as that of the sonic hedgehog (Farmer et al., 2016; Gingrich *et al*., 2022; Xie et al., 2022), necessitates cellular rather than population measures of activity. To expand our understanding of the relationship between astrocyte calcium dynamics and animal behavior, we developed a high-resolution, long working distance, multi-core fiber optic imaging system capable of resolving GCaMP fluorescence changes induced by fluctuations in intracellular calcium within cortical astrocytes in freely behaving mice. By implementing this imaging modality in different behavioral contexts, we show that cortical astrocytes exhibit remarkably diverse calcium activity patterns, which are modulated by internal states and exhibit features appropriate for integrating state transitions over the circadian cycle. This expanded knowledge of the astrocyte behavior-activity framework and the development of technology to explore these interactions will help to define the molecular mechanisms that control astrocyte activity and the consequences of distinct activity patterns on neuronal activity.

### Multi-core fiber optic imaging enables longitudinal monitoring of astrocyte activity *in vivo*

Multi-core optical fiber bundles are frequently used to deliver light to activate optogenetic effectors and monitor fluorescence changes using photometry (Szabo et al., 2014; Tsunematsu et al., 2021; Zhang *et al*., 2021). In detection mode, such systems necessarily integrate light from various sources, providing an ensemble average of population activity dictated by the expression of the sensor, the scattering properties of the tissue, and the photon capture capabilities of the optical path. Coherent multi-core fiber bundles use many thousands of optically isolated fibers to channel light to the detector, preserving spatial information. Here, we developed an objective lens assembled from two pairs of aspheric miniature lenses to collect and focus emitted light onto the end of a 30,000-core fiber bundle to enable cellular resolution of active regions. Importantly, by using lenses with an extended focal length, this removable, lightweight, small form factor device enabled longitudinal imaging through an implanted cranial window with minimal behavioral interruption, providing sufficient spatiotemporal resolution and sensitivity to resolve cellular fluorescence changes within the intact brain while animals were engaged in normative behaviors. Although this system operates in an epifluorescence mode, each optical fiber core intrinsically acts as a spatial filter to prevent photons from outside the focal plane to infiltrate, providing better optical sectioning than a traditional wide-field epifluorescence microscope (Accanto et al., 2022; Sivankutty et al., 2016). The ability to modify imaging parameters, such as light sources, filters, and optic sensors remotely using standard tabletop optical components in this system affords rapid and extensive adaptation to experimental needs. Further modifications to this system could be implemented to enable deeper structure observation and high resolution with multi-photon excitation and/or adaptive optics; fast three-dimensional imaging using adaptive lenses and laser scanning; multicolor detection and laser speckle contrast imaging to resolve interactions between cell types and blood flow; and holographic or random-access fiber illumination to manipulate cellular activity patterns using optogenetics (Accanto *et al*., 2022; Cha *et al*., 2015; Pak *et al*., 2021; Plöschner et al., 2015; Soleimanzad et al., 2019). Although requiring fewer miniature elements than head-mounted miniscopes, this system requires assembly of the multi-lens array and fiber optic coupling. Additionally, although animals acted similarly when tethered or untethered, the multi-core fiber is more rigid than electrical wires used for data transfer in miniscopes, which could potentially influence more subtle behavioral aspects.

### Astrocytes exhibit distinct calcium activity patterns during diversive behaviors

Among other behaviorally relevant state changes, astrocytes have been implicated in controlling sleep homeostasis in multiple species (Blum *et al*., 2021; Clasadonte *et al*., 2017; Ingiosi and Frank, 2022). Astrocyte calcium levels are thought to be the primary means through which these cells accumulate information about the circadian cycle, a hypothesis supported by pioneering imaging studies in unrestrained mice using head-mounted miniscopes (Ingiosi *et al*., 2020; Kjaerby *et al*., 2020; Shekhtmeyster et al., 2021). By implementing our multi-core optical fiber imaging system, we were able to extend this analysis over 24hr imaging periods to resolve longitudinal activity patterns of individual astrocytes within the cerebral cortex, providing direct evidence that calcium activity within individual astrocytes varies prominently with the internal clock of the animals. These findings suggest that astrocytes, in addition to encoding episodic sleep-wake state transitions (Bojarskaite et al., 2020; Ingiosi *et al*., 2020; Kjaerby *et al*., 2020; Thrane *et al*., 2012), can also store an extended record of sleep required for sleep homeostasis (Halassa et al., 2009; Ingiosi and Frank, 2022) through activation of calcium-dependent changes in gene expression (Farhy-Tselnicker *et al*., 2021; Hasel et al., 2017; Kaczmarczyk et al., 2021; Yu et al., 2021).

Using simultaneous astrocyte calcium imaging and behavioral tracking, we linked astrocyte calcium changes to other distinct behaviors using a semi-automated behavior classifier (Figure 4). Consistent with studies in head-immobilized animals (Paukert *et al*., 2014; Slezak et al., 2019), we observed that Q-A transitions were associated with global, transient activation of cortical astrocytes. In contrast, there was a difference among self-paced behaviors, specifically in exploration *vs* maintenance behaviors, with only the former associated with a calcium rise. This differential astrocyte response to behaviors that share similar engagement of motor programs argues against ongoing movement/motor/muscular information processing by cortical astrocytes, a conclusion also drawn from observing animals performing Q-A transitions (Peters et al., 2017). It is notable that, during maintenance behaviors, astrocyte activity fell below the pre-behavioral value (Figure 4C & 8), suggesting that there may be bidirectional modulation of cortical astrocytes, either from active suppression (Mariotti et al., 2018) or from a decrease in neuromodulatory tone, perhaps reflecting a general decrease in noradrenergic or cholinergic neuron firing during satiety (Slezak *et al*., 2019).

To explore with which behavioral features astrocyte activities are most closely associated, we adopted paradigms where mice were allowed to explore arenas and investigate novel objects. These sets of experiments provided behavioral distinctions between exploration and non-exploration. Notably, in paradigms of diversive exploration, it was the behavioral termination that dominated ensemble astrocyte calcium elevations (Figure 5) (Sołyga and Barkat, 2019).Offset-responsive astrocytes might therefore reflect internal adjustments, such as deviations of expected input from contextual feedback. This feature is of particular relevance for generating appropriate behavioral responses to deviations from expected information. Consistent with this conclusion, primary cortices in rodents have been shown to not only represent primitive inputs/outputs, but also act as feedback detectors, essential for rapid initiation of behavioral adaption to the unexpected (Attinger et al., 2017; Lopes et al., 2017). Indeed, recent studies indicate that new inferences of uncertainty are made based on local cholinergic and noradrenergic suppression of intracortical transmission (Vecoven et al., 2020).

Directed exploration paradigms provide unambiguous discrimination of investigatory behaviors and provide an opportunity to observe specific responses of individual cortical astrocytes (Figure 6). Consistent patterns of cortical astrocyte activity across object interactions were reminiscent of the attractor network underlying internal states (Katori et al., 2011). Within a given group of astrocytes, the differential timing of single cells in sub-clusters suggests context-dependent modulation of individual cells. We conclude that, together with known changes of ensemble activity that serve as a basic structure, the staggered recruitment of astrocytes could prolong neuromodulatory events and provide a means to encode distinct temporal sequences of neuronal activity.

Intriguingly, during exploration directed to novel objects, astrocytes exhibited a trial-by-trial reduction in calcium transient magnitude. This fast adaptation implies that the responsiveness of cortical astrocytes, on a population level, was updated rapidly when animals were in a familiar context. By suppressing the responses to experienced contexts, cortical astrocytes may gain the capacity to encode contingencies more efficiently and to maintain responsiveness to rare saliencies (Levy and Bargmann, 2020). However, within a population there were always astrocytes that did not exhibit the principal features of defined clusters. These cells were either unresponsive throughout or presented unreliable patterns across trials. As in the case of fast-adapting ensembles, cortical astrocytes that fell out of clusters in this novel object-exploring context could serve as a reserve to permit circuits to encode a broad range of salient features (Moeyaert et al., 2018).

### Astrocytes integrate input from multiple neuromodulators during flexible behaviors

To investigate the molecular mechanisms that enable engagement of cortical astrocytes in different contexts (Figure 7), we performed local pharmacological manipulations to restrict receptor engagement to superficial cortical areas. As shown previously, NE induced consistent increases in calcium throughout the imaging field during Q-A transitions through activation of α1 adrenoceptors (Paukert *et al*., 2014; Srinivasan *et al*., 2015; Thrane *et al*., 2012). Moreover, NE and ACh inputs converged and enabled synergistic activation during diversive exploration. Mechanistically, once receiving convergent or divergent inputs, individual astrocytes may interpret and integrate various time-dependencies via cross-talk between G-protein-coupled receptors. For example, Gαi/o- and Gαq-linked G-protein-coupled receptors have been shown to yield synergistic calcium responses in the hippocampus of young animals (Durkee et al., 2019) and in astrocytes of developing mice (Kellner et al., 2021). Such interactions could allow more diverse outcomes, as recent studies indicate that chemogenetic activation of Gαi/o or Gαq pathways in astrocytes induced distinct changes in non-rapid eye movement, sleep depth and duration (Vaidyanathan et al., 2021).

In contrast to the NE-dominant response during Q-A transitions, ACh influenced the amplitude of astrocyte calcium transients during diversive explorations, while NE shaped the time course and timing relative to the behavioral sequence. This convergent action may reflect the ambiguous state change involved not only in exploring, but also in confronting the unexpected. Consistent with this possibility, when animals directed their exploration to an unambiguous target, mAChR signaling became more prominent. Using a new genetically encoded optical sensor for ACh (iAChSnFR) with hundred-millisecond kinetics, we found that this ACh reaches astrocyte membranes when mice explored novel objects (Borden *et al*., 2020). A direct action of ACh on astrocytes is also supported by the results of *in vivo* topically applied antagonists in the context of minimal endogenous neural and synaptic activity (Navarrete et al., 2012; Takata et al., 2011). Our studies also suggest that α1 adrenoceptors and group I mGluR signaling have contrasting effects on cell synchrony, with the former enhancing astrocyte network synchrony and the latter promoting desynchrony. This glutamatergic network desynchronization implies that modulation of astrocytes by local neuronal activity or primary sensory inputs, while subtle, does exist (Poskanzer and Yuste, 2016). This is significant because direct engagement of these inputs *in vivo* does not, by itself, promote the widespread activation of astrocytes comparable to that induced by NE during periods of heightened arousal (Paukert *et al*., 2014; Reitman *et al*., 2023; Slezak *et al*., 2019). Thus, different neuromodulators released at specific times during a behavioral sequence appear to converge on astrocytes to manipulate distinct aspects of their calcium activity, suggesting that this is a highly nuanced signaling mode that varies both in space and time depending on which neuromodulators are present.

### Astrocyte calcium activity is modulated by internal state

Many behaviors are strongly modified by internal state. We also explored whether changes in motivation alter the response of astrocytes, focusing on maintenance behaviors (drinking, feeding), which were shown to elicit no activation in cells in animals provided unlimited access to food and water. If cortical astrocytes encode salient contexts, manipulating the internal state of animals without changing the external environment should also enforce state-dependent activation of these cells (Augustine et al., 2019; Burgess et al., 2016). In keeping with this prediction, perturbed homeostasis, like food and water restriction, reversed astrocyte calcium responses from decreasing to profoundly increasing (Figure 8). We further determined that exposing animals to food, but preventing consumption, initiated responses that were even stronger than if they were allowed to eat; the strongest response was observed in cases where mice were able to smell the resource but could not see or access it. This phenomenon further strengthens the conclusion that cortical astrocyte activity is aligned with contextual saliencies, likely reflecting differences in the level of neuromodulatory engagement when goals are predicted but not achieved.

Despite many advances in modeling neural activity in complex networks, astrocytes have been characteristically excluded, since much less is known about their activity patterns within specific behavioral contexts or the effects of this activity on surrounding cells. Recent studies have attempted to address this question by activating or silencing astrocytes within circuits.

However, interpretation of these findings may be complicated by limitations in our ability to mimic the normal spatio-temporal characteristics of their activity or to block these dynamic calcium changes transiently during specific, appropriate behavioral sequences. This challenge may explain why manipulations of astrocytes using standard opto- or chemogenetic approaches, have yielded conflicting results (Chen et al., 2016; Lyon and Allen, 2022; Poskanzer and Yuste, 2016; Sweeney et al., 2016; Yang et al., 2015). New tools are being developed to manipulate calcium signaling in astrocytes, such as the expression of a plasma membrane calcium ATPase to suppress calcium transients (Yu *et al*., 2021), melanopsin to induce calcium activity (Mederos et al., 2019), and long-wavelength opsins to allow modulation of astrocytes in deeper brain regions without fiber or lens insertion (Chen et al., 2021), which may further extend our understanding about the consequences of astrocyte calcium dynamics during different behaviors. The profound consequences of these manipulations raise the intriguing possibility that dysregulation of astrocyte calcium signaling contributes to neuropsychiatric diseases (Carayol et al., 2011; Cederlöf et al., 2015; Pchitskaya et al., 2018). *In vivo* analyses, such as those enabled by this fiber optic imaging platform, will help reveal the precise nature of astrocyte activity in cortical circuits and provide a physiological framework necessary to define their contributions to specific behaviors.

## Acknowledgements

We thank T. Shelly for machining expertise and members of the Bergles Lab for discussions and comments on the manuscript. These studies were supported by grants from the National Institutes of Health (NS050274; MH100024; and MH083728) and a Discovery Award from The Johns Hopkins University.

## Author Contributions

Conceptualization, Y.T.A.G. and D.E.B.; methodology, Y.T.A.G., J.C., R.P. and J.U.K.; investigation, Y.T.A.G. and E.H.; providing reagents, L.L.L. and D.E.B.; formal analysis, Y.T.A.G and E.H.; writing – original draft, Y.T.A.G. and D.E.B.; writing – review and editing, Y.T.A.G., E.H., J.C., R.P., L.L.L., J.U.K. and D.E.B.; funding acquisition, J.U.K. and D.E.B.

## Methods

### Resource availability

#### Lead contact

Requests for sharing resources and reagents should be directed to the Lead Contact, Dwight Bergles.

#### Materials availability

The original imaging system developed here can be accessed upon request made to the lead contact. The details of system assembly were delineated in the paper. iAChSnFR and iAChSnFR-Null constructs and AAV virus are available from Addgene (137950–137962).

#### Data and code availability

All images reported in this paper will be shared by the lead contact upon request. The original code has been deposited and publicly available on GitHub as of the date of publication, with the DOI listed in the key resources table. Any additional information required to reanalyze the data reported in this paper is available from the lead contact upon request.

### Experimental model and subject details

#### Subject

We included in this paper mice of random sex, mixed *B6N;129* background (from genetically-modified animal crosses as below) and between 20 to 24-weeks-old. The animals could assess water and food *ad libitum* in standard polycarbonate cages with enrichment. The housing facilities are maintained at 40–60% humidity, at a temperature of 20–25°C and on a 12-hour light/dark cycle. All experiments and procedures were approved by the Johns Hopkins Institutional Care and Use Committee.

#### Transgenic models

Generation of the following mouse lines have been previously published: *Tg(Slc1a3-cre/ERT)1Nat/J*, also known as *GLAST-CreER* (Wang et al., 2012); STOCK *Gt(ROSA)26Sor^tm1.1(CAG-EGFP)Fsh^/Mmjax*, also known as *RCE:loxP* (Sousa et al., 2009), *B6N;129-Gt(ROSA)26Sor^tm1(CAG-GCaMP3)Dbe^/J*, also known as *Rosa26-lsl-GCaMP3* (Paukert *et al*., 2014). Corresponding RRIDs are listed in key resources table. We acquired these mice from the Jackson Laboratory and crossed the Cre recombinase-conditional EGFP (*RCE:loxP*) or GCaMP (*Rosa26-lsl-GCaMP3*) to the Cre-bearing (*GLAST-CreER*) mice. Offspring (*GLAST-CreER; RCE:loxP* or *GLAST-CreER; Rosa26-lsl-GCaMP3* mice) were then exposed to tamoxifen to induce selective expression of EGFP or GCaMP in astrocytes.

### Method Details

#### Tamoxifen Injections

The transgenic mice received 3 intraperitoneal injections of 100 mg/kg body weight tamoxifen (T5648-1G, MilliporeSigma, CAS:10540-29-1), dissolved in sunflower seed oil (S5007, MilliporeSigma, CAS: 8001-21-6), within 5 days during the fourth postnatal week.

#### Cranial window implantation

The 20 week-old mice were anesthetized using isoflurane (Baxter International) flowed in O_2_ (4% in 1 l/min for induction and 1.5% in 0.5 l/min for maintenance, targeting at 1-2 breaths per second). The central temperature was kept at 37.5 ℃ with a feed-back controller (TC-1000, CWE Inc). We adopted peri-operative occlusion and lubricating eye-ointments (GenTeal PM) to prevent ocular complications. After thorough shaving and skin asepsis using 3 alternative swabs of 70% Ethanol & 10% Povidone-Iodine, a 10 mm-long rostrocaudal opening, revealing landmark sutures, was made in the scalp. Then, subcutaneous tissue and muscles were trimmed and periosteum removed. Tissue margins were secured with super glue (KG 483, Krazy Glue). A stainless steel headplate with a central opening was cemented to the exposed skull (C&B Metabond, Parkell; 1520 BLK, Lang Dental). Animals were then positioned to a head-fixed apparatus. Using a hand-piece micro-drill (XL-230, Osada) drilling beyond the cancellous bone, followed by custom ophthalmic blade (Superior Platinum, ASTRA; 14134, World Precision Instrument) cutting the inner compact bone, a craniotomy (2.8 mm x 3.0 mm) was made at 1.0 mm anterior to lambda and 3.1 mm lateral from midline. A custom single-edge rounded trapezoidal glass coverslip (85-130 µm in thickness) was thereafter placed and cemented in place (C&B Metabond, Parkell; 1520 BLK, Lang Dental).

#### In vivo imaging

A 0.75 W, 473 nm, 0.8 mm beam of an optically pumped semiconductor laser (OBIS473LX, Coherent) was expanded (Olympus PLN 20×/0.4) and reflected by a dichroic mirror (FF499-Di01-25x36, Semrock). The expanded and reflected beam was then coupled into a multi-core optical fiber bundle (FIGH-30-650S, Fujikura) *via* a focusing objective (Olympus PLN 10×/0.25). At the distal fiber end, light was delivered to tissue through a 470-570 nm achromatic pair of aspheric doublets (352140-A, Thorlabs) housed in stainless steel tubes (Small Parts). Returning light travelled back *via* the lens probe, the optical fiber, the coupling objective, the dichroic mirror, an emission filter (MF525-39, Thorlabs), a fixed lens (AC254-150-ML-A, Thorlabs) and was focused onto a CCD (GS2-FW-14S5M-C, FLIR).

For the reflectance imaging setup, dual spectral reflectance at major excitation & emission wavelengths of GCaMP fluorescence imaging were sampled as reported (Bouchard et al., 2009). Using a long-pass dichroic (FF499-Di01-25x36, Semrock), we combined cyan (OBIS473LX - measured peak λ: 473 nm -, Coherent) and green (PLP520 - measured peak λ: 523 nm -, Osram) illumination into one path, further coupled into the fiberscope described in the section above. Using a beamsplitter (EBS1, Thorlabs), reflectance signals were directed into the detecting path, and images were acquired with a single 16-bit depth scientific complementary metal-oxide-semiconductor camera (C11440-42U30, Hamamatsu Photonics).

We exploited a motorized micro-manipulator (MX7630L, DR1000, MC1000e, Siskiyou; GNL10, Thorlabs; custom machine parts) for precision positioning of the fiberscope to a desired FOV; mini-objective mounts were nutted down on headplates to allow optics-animal coupling; FOV was then secured using screws to hold the mini-objective in place (Figure 2A). Data were acquired at 20 frames per second, on-line averaged to 2 or 4 frames per second, with a resolution of 1280 × 960 pixels using commercial software development kit (FlyCapture SDK, FLIR) integrated into the master program (C#, Microsoft).

Two-photon head-fixed imaging was conducted using a laser-scanning microscope (LSM 710 NLO, Carl Zeiss) equipped with a long working distance objective (W Plan-Apochromat 20x/1.0 DIC CG 0.17). With a 920 nm output from a Ti:Sapphire laser (Chameleon Ultra II, Coherent), 250 μm by 250 μm images of two-photon excited GCaMP fluorescence were acquired at 512 by 512 pixel resolution. The acquisition rate was four frames per second.

#### Quantitative assessment of behaviors

The imaging suite was set up in an isolated room so as to limit potential stressors. Ceiling-reflected incandescent lights at corners of the room provided lighting measured 28-32 lux (0.04 W/m^2^, 555 nm) in the center of mice behaving area. Ambient sound was masked by a white noise machine (BRRC112, Big Red Rooster). After recovering from window implantation, mice bore weights on their heads (3.0 g) in cages for two weeks; during the same period of time, we habituated mice to experimental settings. Experiments were carried out in the fourth week, and each animal would engage in only one single paradigm unless otherwise mentioned. Mice lived on a 12-hour light/dark cycle, and were subjected to experiment during their dark, active phase (between 18:00 and 06:00). We transported mice to the suite 1 hour prior to testing, and during experiment, stayed the unnoticeable to them.

Two near infra-red cameras (NIRCam) (DCC1645C, Thorlabs – with IR filter removed, or FI8910W, Foscam) were centered 100 cm above (NIRCam-1), and on the same level 25 cm (NIRCam-2) from the experimental platform. Videos were recorded at 30 frames per second with a resolution of 1280 × 960 pixels through master program capture of video streams (Foscam VMS, Foscam; C#, Microsoft).

For the head-fixed quiet awake to active awake (Q-A) paradigm, mice were placed on a custom head-fixed rotating platform (S1-Round12-.125, Source One), set on the stepper motor-driven mode (ROB-09238, ROB-12779, SparkFun Electronics). Rotation was monitored with an optical encoder (600128CBL, Honeywell). An Arduino Mega 2560 R3 microcontroller board (191, Adafruit) signaled motor onsets and digitized encoder readings.

To assess the influence of fiberscope-tethering on animals, mice explored a 30 x 30 x 30 cm^3^ chamber for 30 minutes. Then we recorded locomotion for the following 30 minutes. A second group of mice went through the same protocol but without being tethered to the fiberscope. Before testing and between mice, the chamber was cleaned with 3 alternative washes of purified H_2_O & 70% ethanol.

For longitudinal behavioral cycle imaging, mice were habituated to the imaging chamber for increasing durations beginning two weeks prior to the imaging day. In the three days prior to the imaging day, mice were tethered to the fiberscope and allowed to freely move for 60 minutes each day. On imaging day, mice were tethered and freely moving in the imaging cage for 60 minutes prior to the start of data collection. Mice were then allowed to explore the chamber, with the experimenter monitoring the mice for stress or discomfort remotely. Recordings were terminated after 26 hrs, after which mice were untethered and returned to their home cage. To avoid potential confounds from handling and stress, the first two hrs of data collection were discarded.

In order to phenotype spontaneous behavior of mice in home cages, we separated and housed individual mice in single cages 26–28 hrs prior to imaging day. On the day of experiment, we performed imaging when each mouse behaved freely in its home cage. During imaging, an unfamiliar object to the mouse was placed in the cage to stimulate the exploratory and investigatory aspects of behaviors, which animal would infrequently do at home. After experiments, mice were re-introduced with their previous cage mates under supervision.

To observe diversive exploration in environmental novelty, mice were placed in the center of a 30 x 30 x 30 cm^3^ chamber and imaged for 5 minutes. Upon each mouse completing a trial, we placed it into a new, interim cage until all his or her cage mates have been tested. Then mice from the same original cage returned to their home. A separate group of mice was allowed to explore the chamber for 60 minutes 24 hrs before. They were returned to the home cage afterwards, and then participated in the same paradigm as the experimental animals on the day of imaging. Before testing and between mice, we cleaned the chamber with 3 alternative washes of purified H_2_O & 70% Ethanol.

For inspective novelty exploration, on the day before imaging, mice got habituated to a 40 x 20 x 30 cm^3^ chamber for 60 minutes. On imaging day, mice first interacted and became familiar with two identical objects (positioned at two interior corners of chamber, 5 cm from walls) for 10 minutes. Afterwards mice stayed in an interim cage, during which we replaced the two objects, one with an identical triplicate and the other with a novel object. Within an hour, mice returned to the testing chamber and explored the familiar *vs.* the novel object for 10 minutes at free will. We used 250 ml polypropylene copolymer jars (about 6.4 cm in diameter and 11.5 cm in height) of light blue desiccants (2118-0008, Thermo Fisher Nalgene; 23001, Drierite), and towers of polyethylene-wrapped colored polyhedron sitting on color-taped optical posts (RS3P4M, Thorlab) (about 5.0 cm in diameter and 14.0 cm in height) as the target objects. Choice of objects for familiar and novel trials were based on a crossover design to avoid confounding due to differences in objects. Before testing, between sessions and between mice, we cleaned the chamber with 3 alternative washes of purified H_2_O & 70% Ethanol.

Laser illumination, brain imaging and behavior recording were synchronized on a generic data acquisition board (USB 6009, National Instruments). When behavioral recording cannot be triggered on the master clock, the frame clocks of the CCD were duplicated to set on a 940 nm light emitting diode (H&PC-56931, Chanzon), recorded in NIRCam recordings as timestamps; local time at millisecond precision was also tagged in NIRCam images (DateTime.Now.ToString; C#). We programed the master control on the basis of off-the-shelf development kits (FlyCapture SDK, FLIR; Foscam VMS, Foscam; Measurement Studio Standard, National Instruments; C#, Microsoft) running in a standard x64 operating system (Windows 7 Enterprise, Microsoft), on a custom-assembled desktop computer (MZ-7KE256BW, Samsung; BX80648I75930K, Intel; CMK32GX4M2B3200C16, CORSAIR; STRIX-GTX960-DC2OC-2GD5, Asus; GA-X99-UD3P, GIGABYTE).

#### Pharmacology

We performed systemic pharmacology by injecting mice who participated in freely-moving imaging intraperitoneally with 10 ml/kg 0.9% NaCl (vehicle) in control mice or 10 ml/kg 0.9% NaCl with dexmedetomidine (3 µg/kg) in experimental groups.

For topical pharmacology in freely-moving mice, we first fabricated chemical-infusing cannula using methods as previously reported (Kokare et al., 2011). In brief, a guide cannula of 3.0 mm in length was made out of a 23G hypodermic needle (305120, Becton Dickinson). We layered epoxy (20945, Devcon) to a half-circle baseplate - about 1.5 mm in radius - at 1.0 mm from one end, denoted as the intracranial end. Second, a 5.5 mm-long stylet was made from a 30G stainless steel wire (AW1-30-SS, Artistic Wire), inserted and protruding from the intracranial end for 0.1 mm and for 2.4 mm from the other.

Implantation of cannula is enabled in a two-step operation. We first cemented headplates to the skulls of 20-week-old animals, and then delayed the next step of surgery, including implanting windows and cannulas, to the day before imaging in the 24^th^ postnatal week. To provide infusion access to imaging area, two quarter-circles at two diagonally opposite corners -0.35 mm in radius - were cut off from the custom-shaped coverslip. During surgeries, additional skull openings of 0.7 mm in diameter were made at corners of craniotomy matching the quarter-circle cuts of coverslip. Dura was removed. Once coverslip is cemented, we disinfected and filled cannulas with 3 alternative washes of sterile 0.9% NaCl (Baxter Healthcare) & 70% Ethanol, placed stylet-in-guides at two corners, cement-set the baseplate on skull (Grip Cement, Dentsply International), sealed coverslip corners with tissue adhesive (VetBond, 3M) or UV curable optical adhesive (NOA61, Norland Products), and built layers of cement around guides (Grip Cement, Dentsply International). The extracranial end was bent at right angle and covered with silicone (Ecoflex 5, Smooth-On).

On the imaging day, an internal cannula of 5.5 mm in length was made out of the 30G hypodermic needle (305106, Becton Dickinson). We fitted 2.4 mm of internal cannula into a PE10 tube (427401, Becton Dickinson) and sealed with epoxy (20945, Devcon). We disinfected and filled the internal cannula with 3 alternative washes of 70% ethanol & sterile artificial cerebrospinal fluid (aCSF 298 Osm/kg, pH 7.35 adjusted with NaOH), containing in mM: 137 NaCl, 2.5 KCl, 1 MgCl_2_, 2 CaCl_2_, 20 HEPES. For mice presenting desired behaviors in a baseline session, we repositioned them in a head-fix apparatus, removed silicone coverings, replaced their two stylets with internal cannulas, rendering ∼0.1 mm extension below the guides, extensively sealed the extracranial cannula system (20945, Devcon), perfused the cortex at a flow rate of 0.5 ml/min for 15 minutes with drugs (100 μM prazosin, 100 µM atropine, 30/30 µM MTEP/JNJ16259685, 30 μM Methysergide) dissolved in aCSF, and drained the overflow through the other cannula. Then we removed the internal cannulas, sealed the openings with epoxy (20945, Devcon), released the animals into the behavior chambers, and started a session to evaluate pharmacological effects on astrocyte dynamics.

We selected 24-week-old mice for the topical pharmacology during head-fixed two-photon imaging. To perfuse drugs and keep stable imaging, during surgeries we removed dura and covered the 2.5 x 2.5 mm^2^ craniotomy with a double-layered cranial window, which consisted of a 2.0 x 1.5 mm^2^ (130-170 µm thick) glass UV glued (NOA61, Norland Products) to center of a 3.0 x 2.0 mm^2^ (130-170 µm thick). Overhanging edges of the 3.0 x 2.0 mm^2^ were cemented (C&B Metabond, Parkell; 1520 BLK, Lang Dental) to the 3.5 x 3.5 mm^2^ drilled mark, around craniotomy at the depth of inner compact bone. The 2.0 x 1.5 mm^2^ glass hence sat gently on tissue and moderated brain motion. We covered the openings temporarily with silicone (Ecoflex 5, Smooth-On). Experiment took place after mice made full recovery from anesthesia, when we subjected them to a head-fixed awake imaging setting, and flowed drugs at a rate of 5 ml/min (100 μM Carbachol, 100 μM Phenylephrine, 30 μM VU 0255035) dissolved in aCSF containing neurotransmission blockers (1 μM Tetrodotoxin, 100/100 μM NBQX/CPP, 100 μM picrotoxin).

#### Expression of iAChSnFR

The 20 week-old *GLAST-CreER; Rosa26-lsl-GCaMP3* mice were anesthetized and prepped as above. Once revealing landmark sutures, we made a burr hole (1.0 mm in diameter) in the skull using a micro-drill (Stoelting) at 1.0 mm anterior to lambda and 3.1 mm lateral from midline. Adeno-associated virus, *AAV.GFAP.iAChSnFR ^Active^* or *AAV.GFAP.iAChSnFR ^Null^* (Borden *et al*., 2020), was injected 0.45 mm ventral from dura surface. Using a Nanoject II (Drummond Scientific), 132 nl of 5.0e+09 gc AAV was delivered through 10× 13.2 nl injections (pausing 10 s in between) *via* a beveled glass pipette (tip diameter 15-20 µm). After injections, the pipette was removed in steps of 0.1 mm per min, and upon full removal, the burr hole was sealed with tissue adhesive (VetBond, 3M). A cranial window would then be placed.

#### Calcium imaging analysis

We processed imaging data using routines established upon published MATLAB pipelines (R2019b, MathWorks). Shifts in imaging fields were corrected using modified MiN1PIPE hierarchical framework (Lu et al., 2018). Each image series was divided into stable *vs.* unstable segments based on KLT tracking of investigator-validated SURF points in a reference frame (built-in function “estimateGeometricTransform” & system object “vision.PointTracker”); unstable segments were aligned with intensity-based rigid registration (“imregister”); then segments were recombined to form registered series. We then TV-L1 denoised images (Lourakis implementation “TVL1denoise”) (Chambolle and Pock, 2011), and retrieved region of interests (ROIs) using CNMF-E, assuming diameter of the background ring 1.5 timed that of the largest cell (Zhou et al., 2018).

Thereafter, average pixel fluorescence was acquired for each ROI, which was transformed into relative changes in fluorescence, Δ*F/F_0_*. *F_0_* was acquired by kernel density estimate (“ksdensity”). A single image series of n frames were downscaled to n/10 bins, the vector of minimal fluorescence in each bin was calculated, kernel density distributions were estimated through the vector in sliding windows of 25 frames, and then the baseline was approximated to be the modified Akima interpolation (“interp1”) of peak values of the distributions.. Across sessions, ROIs were matched using CellReg, a MATLAB implementation of probabilistic cell tracking (Sheintuch et al., 2017). We identified local maxima (“findpeaks”) with z-scored height > 2.5 as tentative peaks, next confirmed with interactive and iterative fitting (iPeak) (O’Haver). Reflectance measurements were converted into estimates of scattering changes to account for the magnitude of absorbance contamination in both excitation and emission fluorescence bands, in accordance with Beer-Lambert derived formula (Ma *et al*., 2016).

For ratio of Fano factor (rFF) computation, we calculated the relative changes in mean astrocyte response and across-event variance (Montijn *et al*., 2014). The fractional changes in FF from epochs of a given behavioral dimension (*i.e.*, moment before rearing, onset of rearing, offset of rearing) to epochs of maximal mean response, taking place during Q-A transitions.

When calculating the posterior probability of mouse activeness during an astrocyte event, we used the following values deducted from our data. The probability of mouse being active was 0.24. The probability of astrocytes having an event was 0.09. For the actual data, the probability of astrocyte having an event when mouse was active was 0.21 ± 0.03 and for the random process 0.09 ± 0.01. We acquired probability density function with 1000 bootstraps or random generators.

#### Behavior analysis

Home cage behavioral phenotypes were classified using published algorithms (Janelia Automatic Animal Behavior Annotator, JAABA) (Kabra *et al*., 2013) adapted to behavior recognition in side-view videos (vision-based system for automated mouse behavior recognition) (Jhuang *et al*., 2010). We trained behavioral classifiers on a 30-min video recorded on NIRCam-2. To evaluate the classifiers’ performance, we selected several random data segments outside of training data, consisted of 50,000 frames in total. Ground-truth labels were provided by investigators independent of this research with three levels of confidences: important, unimportant an unknown. Inter-annotator confusion rates were low for important frames (10.1%), justifying assessment of classifier performance based on important frames (error rates: 7.1%).

To quantitatively measure individual behaviors, *c*ustom graphical user interfaces (GUIs) for individual behaviors were programmed using MATLAB (R2019b, MathWorks). In assessing locomotion, we read in positions of animals on NIRCam-1 (“Track”), and calculated movement time, animal traces, distance traveled & walking speed. In evaluating diversive exploration, we identified in NIRCam-2 videos mouse nose point & tail base (“Rear”), deducted upright truncal extension (Δh), and estimated rearing onset, offset & duration (Δt). In determining inspective exploration, we computed mouse anterior-posterior axis in NIRCam-1 videos (“Object”), recognized when the axis were oriented toward & the nose were in the interaction zone (from centroid to 2.5 cm-wide perimeter around an object), and placed a flag on the time of object exploration.

#### Immunofluorescence

We intraperitoneally administered mice with a euthanasia dose of pentobarbital (100 mg/kg body weight). Once in deep anesthesia, animals were perfused with phosphate buffer saline (PBS) followed by cold 4% paraformaldehyde (PFA). Brains were harvested, post-fixed in 4% PFA at 4°C overnight, cryoprotected in 30% sucrose and sectioned into 50 μm-thick slices on a freezing microtome (Leica SM 2010R). Free-floating sections were washed in PBS, incubated for 1 hr in blocking solution (0.3% Triton X-100 & 5% NDS), and kept at 4°C overnight with primary antibodies (chicken anti-GFP, Aves Labs, 1:4000; rabbit anti-GFAP, Agilent Pathology Solutions, 1:500; mouse anti-S100B, MilliporeSigma, 1:400; mouse anti-NeuN, MilliporeSigma, 1:400; guinea pig anti-Olig2, Bennett Novitch, 1:20000) in 0.3% Triton X-100 and 5% NDS. On the day after, sections were washed with PBS, incubated for 2 hr at room temperature with secondary antibodies (donkey anti-chicken, Alexa Fluor 488, 1:2000; donkey anti-rabbit, DyLight 650, 1:2,000; donkey anti-mouse, Cy™3, 1:2,000; donkey anti-guinea pig, Cy™3, 1:2000) in 5% NDS, washed again in PBS, incubated with DAPI (1μg/ml), mounted on slides and coverslip sealed with mounting medium (Aqua-Poly/Mount, Polysciences #18606-20). Antibody details are shown in Key Resources Table. Epifluorescence Images were taken using Axio Imager.M1 microscope (Carl Zeiss) with a Plan-Apochromat 20x/0.8 M27 objective (420650-9901, Carl Zeiss). Confocal images were acquired using a Zeiss LSM 510 Meta microscope (Carl Zeiss) with an EC Plan-Neofluar 40x/1.30 Oil DIC M27 (420462-9900, Carl Zeiss) objective. We then denoised images using a two-dimensional 3.0 x 3.0 pixels^2^ Gaussian filter, an integrated class in the open source software Fiji (Schindelin et al., 2012).

### Quantification and statistical analyses

Statistics was performed using R (3.5.1, The R Foundation) and MATLAB (R2019b, MathWorks). We conducted parametric statistics assuming normal distribution and equal variances, except when a significant Shapiro-Wilk test warranted nonparametric statistics. When unequal variances were noted, we accepted results of parametric test, with Brown-Forsythe test reported for the experimental data. In using repeated measures analysis of variance (RM-ANOVA), we applied Greenhouse-Geisser ε correction in the event that Mauchly’s tests suggested violation of sphericity. Figure 2C: 2-sample *t*-test; Figure 2F: two-way mixed rank test, ANOVA-type (nparLD: “f1.ld.f1”) (Noguchi et al., 2012); Figure 2G: two-way repeated measures ANOVA, *post hoc* Sidak’s multiple comparison. Figure 3E: Wilcoxon signed-rank test; Figure 3F 2-sample K-S test. Figure 5H: Wilcoxon signed-rank test. Figure 6A: paired *t*-test; Figure 6G: Friedman test, *post hoc* Dunn’s multiple comparison; Figure 6C: two-way repeated measures ANOVA, *post hoc* Sidak’s multiple comparison; Figure 7A: one-way ANOVA with *post hoc* Sidak’s multiple comparison and Kruskal-Wallis with *post hoc* Dunn’s multiple comparison; Figure 7C: one-way ANOVA, *post hoc* Sidak’s multiple comparison; Figure 7F: mixed-effects model, *post hoc* Sidak’s multiple comparison; Figure 7G: mixed-effects model, *post hoc* Sidak’s multiple comparison; Figure 8C: Wilcoxon rank-sum test and Kruskal-Wallis with *post hoc* Dunn’s multiple comparison were reported as mean ± s.e.m and 95% confidence interval (CI). *P* < 0.05 was considered significant.

## KEY RESOURCES TABLE

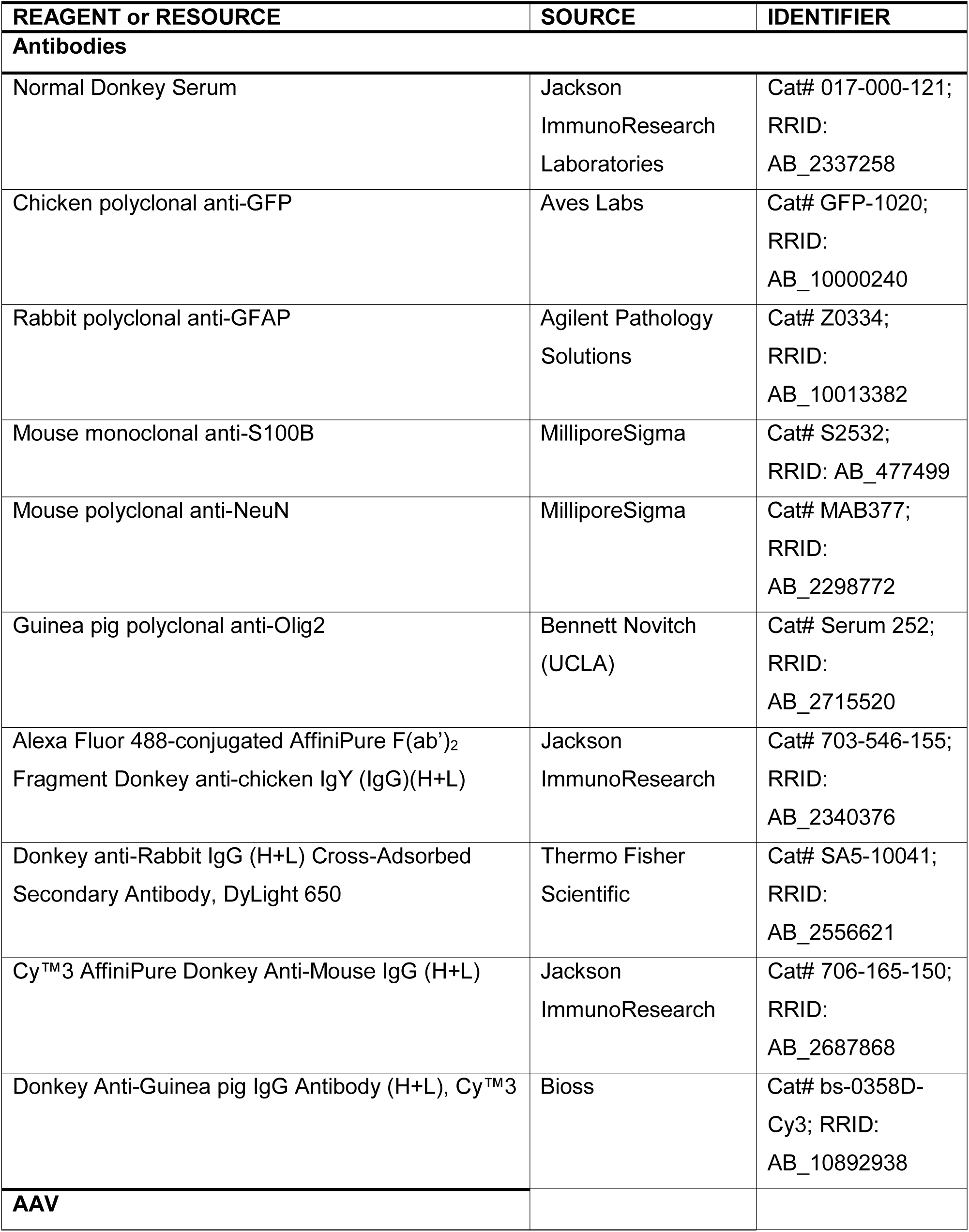
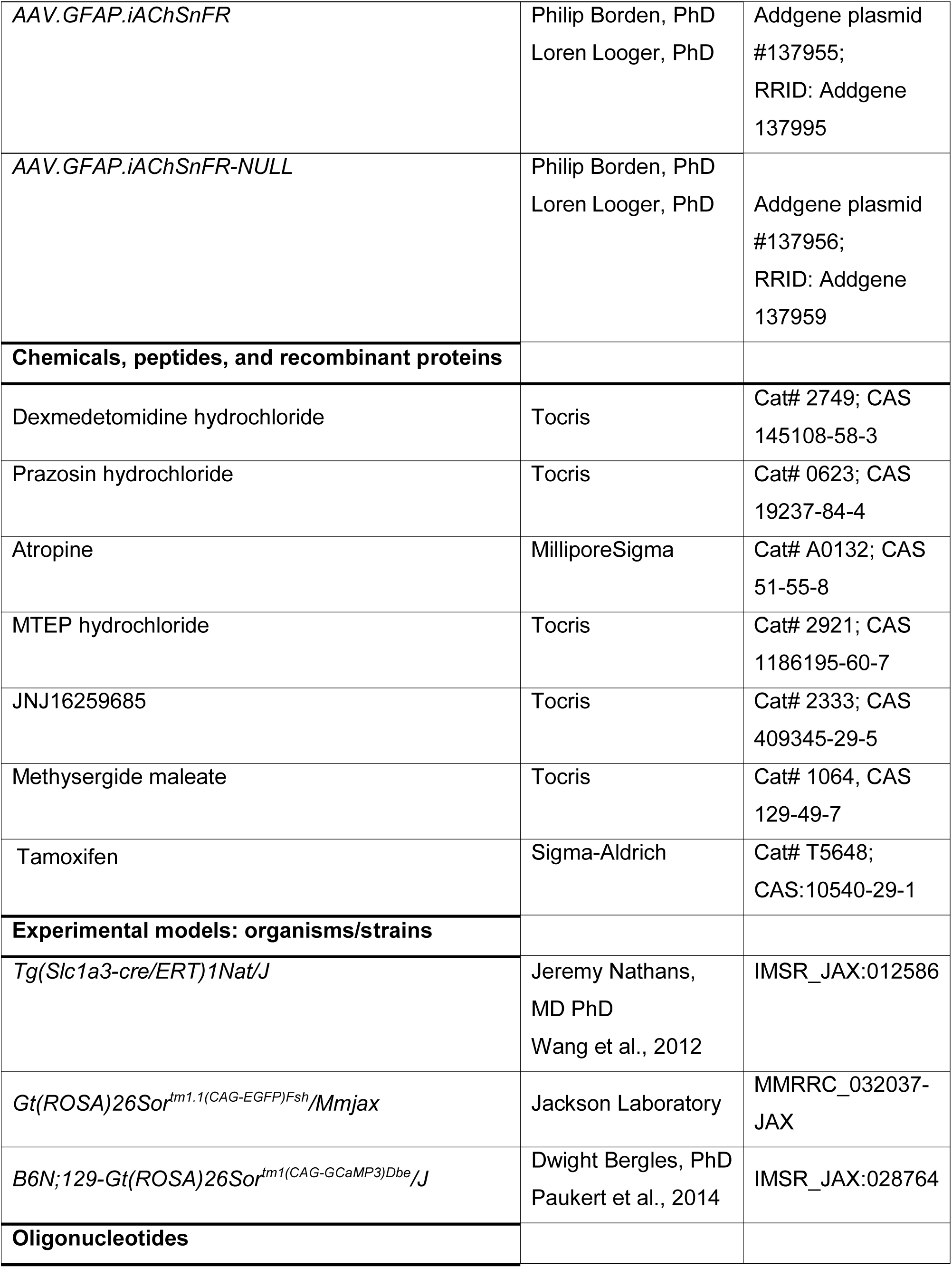
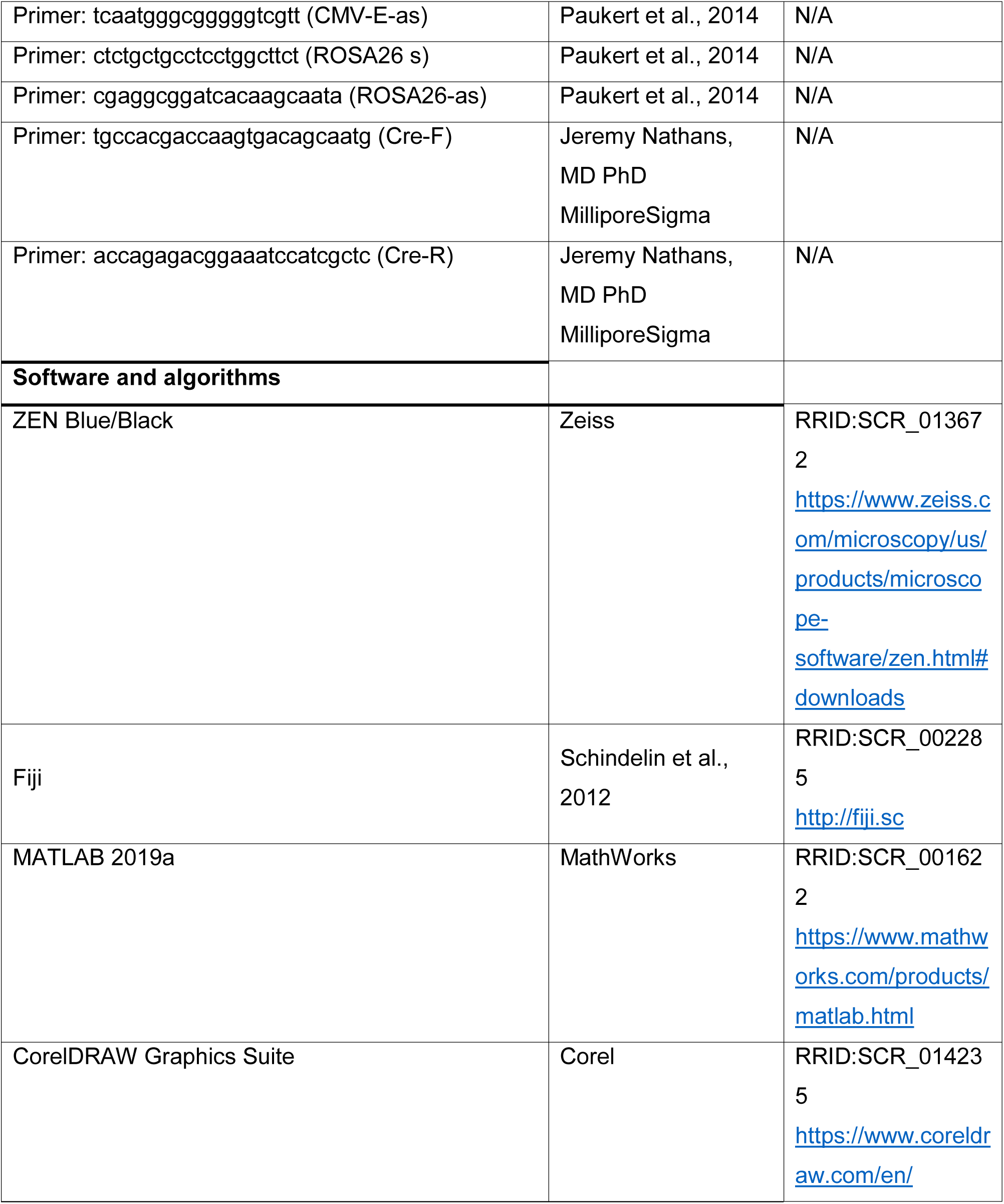
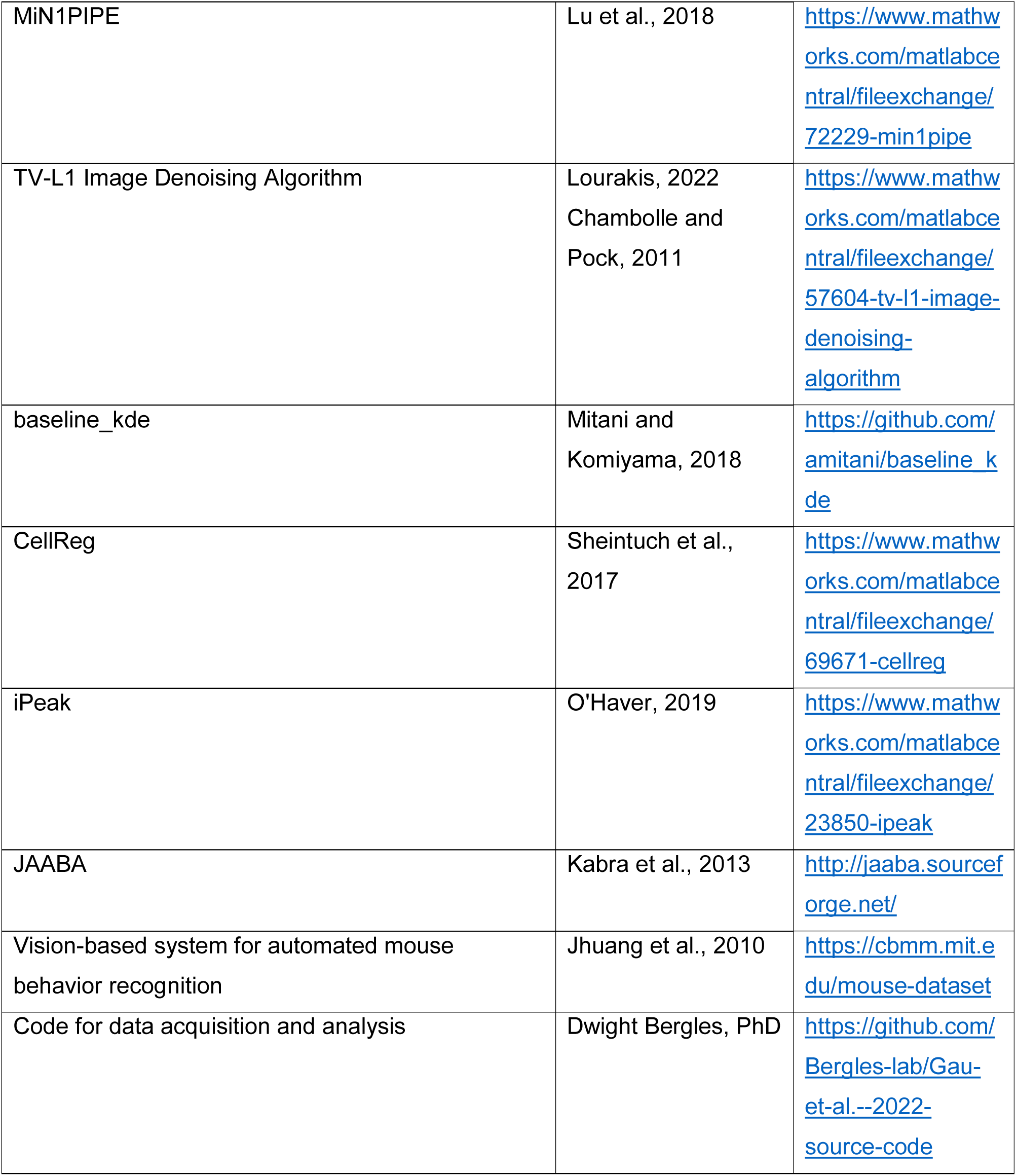

## Supplemental Items

**Figure S1.**
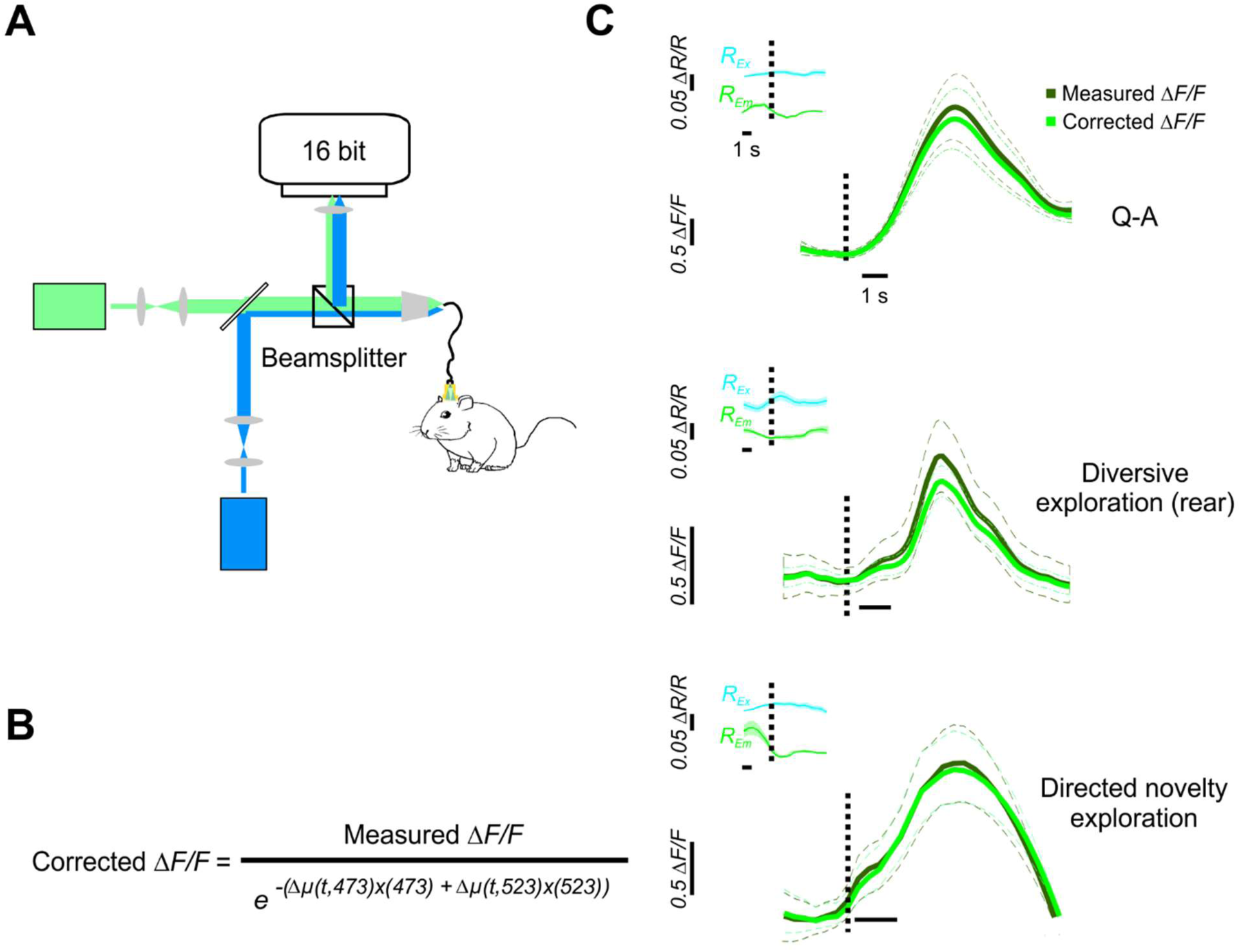
Imaging reflectance change during behaviors. (A) Freely moving reflectance imaging set up at 473 nm and 523 nm, peak excitation/emission wavelengths used for GCaMP imaging. (B) Formula for correcting GCaMP fluorescence signals for associated hemodynamic changes. (C) Reflectance changes had a small effect on the amplitude and time course of GCaMP fluorescence signals in the major behaviors studied in this paper.

## References

Accanto, N., Blot, F.G., Lorca-Cámara, A., Zampini, V., Bui, F., Tourain, C., Badt, N., Katz, O., and Emiliani, V. (2022). A flexible two-photon fiberscope for fast activity imaging and precise optogenetic photostimulation of neurons in freely moving mice. Neuron.

Ackerman, S.D., Perez-Catalan, N.A., Freeman, M.R., and Doe, C.Q. (2021). Astrocytes close a motor circuit critical period. Nature 592, 414–420.

Acquas, E., Wilson, C., and Fibiger, H.C. (1996). Conditioned and unconditioned stimuli increase frontal cortical and hippocampal acetylcholine release: effects of novelty, habituation, and fear. Journal of Neuroscience 16, 3089–3096.

Akdemir, E.S., Woo, J., Huerta, N.A.B., Lozzi, B., Groves, A.K., Harmanci, A.S., and Deneen, B. (2022). Lunatic Fringe-GFP Marks Lamina-Specific Astrocytes That Regulate Sensory Processing. Journal of Neuroscience 42, 567–580.

Attinger, A., Wang, B., and Keller, G.B. (2017). Visuomotor coupling shapes the functional development of mouse visual cortex. Cell 169, 1291–1302. e1214.

Augustine, V., Ebisu, H., Zhao, Y., Lee, S., Ho, B., Mizuno, G.O., Tian, L., and Oka, Y. (2019). Temporally and spatially distinct thirst satiation signals. Neuron 103, 242–249. e244.

Barber, H.M., Ali, M.F., and Kucenas, S. (2021). Glial Patchwork: Oligodendrocyte Progenitor Cells and Astrocytes Blanket the Central Nervous System. Front Cell Neurosci 15, 803057. 10.3389/fncel.2021.803057.

Bayraktar, O.A., Bartels, T., Holmqvist, S., Kleshchevnikov, V., Martirosyan, A., Polioudakis, D., Ben Haim, L., Young, A.M., Batiuk, M.Y., and Prakash, K. (2020). Astrocyte layers in the mammalian cerebral cortex revealed by a single-cell in situ transcriptomic map. Nature neuroscience 23, 500–509.

Bélanger, M., and Magistretti, P.J. (2022). The role of astroglia in neuroprotection. Dialogues in clinical neuroscience.

Berlyne, D.E. (1950). Novelty and curiosity as determinants of exploratory behaviour. British journal of psychology 41, 68.

Blum, I.D., Keleş, M.F., Baz, E.-S., Han, E., Park, K., Luu, S., Issa, H., Brown, M., Ho, M.C., and Tabuchi, M. (2021). Astroglial calcium signaling encodes sleep need in Drosophila. Current Biology 31, 150–162. e157.

Bojarskaite, L., Bjørnstad, D.M., Pettersen, K.H., Cunen, C., Hermansen, G.H., Åbjørsbråten, K.S., Chambers, A.R., Sprengel, R., Vervaeke, K., Tang, W., et al. (2020). Astrocytic Ca(2+) signaling is reduced during sleep and is involved in the regulation of slow wave sleep. Nat Commun 11, 3240. 10.1038/s41467-020-17062-2.

Borden, P.M., Zhang, P., Shivange, A.V., Marvin, J.S., Cichon, J., Dan, C., Podgorski, K., Figueiredo, A., Novak, O., and Tanimoto, M. (2020). A fast genetically encoded fluorescent sensor for faithful in vivo acetylcholine detection in mice, fish, worms and flies.

Bouchard, M.B., Chen, B.R., Burgess, S.A., and Hillman, E.M. (2009). Ultra-fast multispectral optical imaging of cortical oxygenation, blood flow, and intracellular calcium dynamics. Optics express 17, 15670–15678.

Bouvier, D.S., Jones, E.V., Quesseveur, G., Davoli, M.A., A Ferreira, T., Quirion, R., Mechawar, N., and Murai, K.K. (2016). High resolution dissection of reactive glial nets in Alzheimer’s disease. Scientific reports 6, 1–15.

Burgess, C.R., Ramesh, R.N., Sugden, A.U., Levandowski, K.M., Minnig, M.A., Fenselau, H., Lowell, B.B., and Andermann, M.L. (2016). Hunger-dependent enhancement of food cue responses in mouse postrhinal cortex and lateral amygdala. Neuron 91, 1154–1169.

Cabanac, M. (1971). Physiological Role of Pleasure: A stimulus can feel pleasant or unpleasant depending upon its usefulness as determined by internal signals. Science 173, 1103–1107.

Carandini, M., Shimaoka, D., Rossi, L.F., Sato, T.K., Benucci, A., and Knöpfel, T. (2015). Imaging the awake visual cortex with a genetically encoded voltage indicator. Journal of Neuroscience 35, 53–63.

Carayol, J., Sacco, R., Tores, F., Rousseau, F., Lewin, P., Hager, J., and Persico, A.M. (2011). Converging evidence for an association of ATP2B2 allelic variants with autism in male subjects. Biological psychiatry 70, 880–887.

Cederlöf, M., Bergen, S.E., Långström, N., Larsson, H., Boman, M., Craddock, N., Östberg, P., Lundström, S., Sjölander, A., and Nordlind, K. (2015). The association between Darier disease, bipolar disorder, and schizophrenia revisited: a population-based family study. Bipolar disorders 17, 340–344.

Cha, J., and Kang, J.U. (2014). Spatially multiplexed fiber-optic microscopy for simultaneous imaging of multiple brain regions. (Optical Society of America), pp. AF1B. 3.

Cha, J., Kim, D., Cheon, G.W., and Kang, J.U. (2015). Spatially Multiplexed Fiber-optic SLM Microscopy for Applications of Optogenetics. (Optical Society of America), pp. IM4A. 2.

Chai, H., Diaz-Castro, B., Shigetomi, E., Monte, E., Octeau, J.C., Yu, X., Cohn, W., Rajendran, P.S., Vondriska, T.M., and Whitelegge, J.P. (2017). Neural circuit-specialized astrocytes: transcriptomic, proteomic, morphological, and functional evidence. Neuron 95, 531–549. e539.

Chambolle, A., and Pock, T. (2011). A first-order primal-dual algorithm for convex problems with applications to imaging. Journal of mathematical imaging and vision 40, 120–145.

Chaturvedi, R., Stork, T., Yuan, C., Freeman, M.R., and Emery, P. (2022). Astrocytic GABA transporter controls sleep by modulating GABAergic signaling in Drosophila circadian neurons. Current Biology 32, 1895–1908. e1895.

Chen, N., Sugihara, H., Kim, J., Fu, Z., Barak, B., Sur, M., Feng, G., and Han, W. (2016). Direct modulation of GFAP-expressing glia in the arcuate nucleus bi-directionally regulates feeding. Elife 5, e18716.

Chen, N., Sugihara, H., Sharma, J., Perea, G., Petravicz, J., Le, C., and Sur, M. (2012). Nucleus basalis-enabled stimulus-specific plasticity in the visual cortex is mediated by astrocytes. Proceedings of the National Academy of Sciences 109, E2832–E2841.

Chen, R., Gore, F., Nguyen, Q.-A., Ramakrishnan, C., Patel, S., Kim, S.H., Raffiee, M., Kim, Y.S., Hsueh, B., and Krook-Magnusson, E. (2021). Deep brain optogenetics without intracranial surgery. Nature biotechnology 39, 161–164.

Choi, I.S., Kim, J.H., Jeong, J.Y., Lee, M.G., Suk, K., and Jang, I.S. (2022). Astrocyte-derived adenosine excites sleep-promoting neurons in the ventrolateral preoptic nucleus: Astrocyte-neuron interactions in the regulation of sleep. Glia 70, 1864–1885.

Clasadonte, J., Scemes, E., Wang, Z., Boison, D., and Haydon, P.G. (2017). Connexin 43-mediated astroglial metabolic networks contribute to the regulation of the sleep-wake cycle. Neuron 95, 1365–1380. e1365.

Corkrum, M., Covelo, A., Lines, J., Bellocchio, L., Pisansky, M., Loke, K., Quintana, R., Rothwell, P.E., Lujan, R., and Marsicano, G. (2020). Dopamine-evoked synaptic regulation in the nucleus accumbens requires astrocyte activity. Neuron 105, 1036–1047. e1035.

D’yakonova, V. (2014). Neurotransmitter mechanisms of context-dependent behavior. Neuroscience and Behavioral Physiology 44, 256–267.

Devor, A., Hillman, E.M., Tian, P., Waeber, C., Teng, I.C., Ruvinskaya, L., Shalinsky, M.H., Zhu, H., Haslinger, R.H., and Narayanan, S.N. (2008). Stimulus-induced changes in blood flow and 2-deoxyglucose uptake dissociate in ipsilateral somatosensory cortex. Journal of Neuroscience 28, 14347–14357.

Ding, F., O’Donnell, J., Thrane, A.S., Zeppenfeld, D., Kang, H., Xie, L., Wang, F., and Nedergaard, M. (2013). α1-Adrenergic receptors mediate coordinated Ca2+ signaling of cortical astrocytes in awake, behaving mice. Cell calcium 54, 387–394.

Doron, A., Rubin, A., Benmelech-Chovav, A., Benaim, N., Carmi, T., Refaeli, R., Novick, N., Kreisel, T., Ziv, Y., and Goshen, I. (2022). Hippocampal astrocytes encode reward location. Nature 609, 772–778.

Durkee, C.A., Covelo, A., Lines, J., Kofuji, P., Aguilar, J., and Araque, A. (2019). Gi/o protein-coupled receptors inhibit neurons but activate astrocytes and stimulate gliotransmission. Glia 67, 1076–1093.

Endo, F., Kasai, A., Soto, J.S., Yu, X., Qu, Z., Hashimoto, H., Gradinaru, V., Kawaguchi, R., and Khakh, B.S. (2022). Molecular basis of astrocyte diversity and morphology across the CNS in health and disease. Science 378, eadc9020.

Escartin, C., Galea, E., Lakatos, A., O’Callaghan, J.P., Petzold, G.C., Serrano-Pozo, A., Steinhäuser, C., Volterra, A., Carmignoto, G., Agarwal, A., et al. (2021). Reactive astrocyte nomenclature, definitions, and future directions. Nat Neurosci 24, 312–325. 10.1038/s41593-020-00783-4.

Farhy-Tselnicker, I., Boisvert, M.M., Liu, H., Dowling, C., Erikson, G.A., Blanco-Suarez, E., Farhy, C., Shokhirev, M.N., Ecker, J.R., and Allen, N.J. (2021). Activity-dependent modulation of synapse-regulating genes in astrocytes. Elife 10, e70514.

Farmer, W.T., Abrahamsson, T., Chierzi, S., Lui, C., Zaelzer, C., Jones, E.V., Bally, B.P., Chen, G.G., Théroux, J.-F., and Peng, J. (2016). Neurons diversify astrocytes in the adult brain through sonic hedgehog signaling. Science 351, 849–854.

Ferezou, I., Bolea, S., and Petersen, C.C. (2006). Visualizing the cortical representation of whisker touch: voltage-sensitive dye imaging in freely moving mice. Neuron 50, 617–629.

Flusberg, B.A., Nimmerjahn, A., Cocker, E.D., Mukamel, E.A., Barretto, R.P., Ko, T.H., Burns, L.D., Jung, J.C., and Schnitzer, M.J. (2008). High-speed, miniaturized fluorescence microscopy in freely moving mice. Nature methods 5, 935–938.

Gee, J.M., Gibbons, M.B., Taheri, M., Palumbos, S., Morris, S.C., Smeal, R.M., Flynn, K.F., Economo, M.N., Cizek, C.G., and Capecchi, M.R. (2015). Imaging activity in astrocytes and neurons with genetically encoded calcium indicators following in utero electroporation. Frontiers in Molecular Neuroscience 8, 10.

Gelegen, C., Gent, T.C., Ferretti, V., Zhang, Z., Yustos, R., Lan, F., Yang, Q., Overington, D.W., Vyssotski, A.L., and van Lith, H.A. (2014). Staying awake–a genetic region that hinders α2 adrenergic receptor agonist-induced sleep. European Journal of Neuroscience 40, 2311–2319.

Ghosh, K.K., Burns, L.D., Cocker, E.D., Nimmerjahn, A., Ziv, Y., El Gamal, A., and Schnitzer, M.J. (2011). Miniaturized integration of a fluorescence microscope. Nature methods 8, 871–878.

Gingrich, E.C., Case, K., and Garcia, A.D.R. (2022). A subpopulation of astrocyte progenitors defined by Sonic hedgehog signaling. Neural Development 17, 1–13.

Giovannini, M.G., Bartolini, L., Kopf, S.R., and Pepeu, G. (1998). Acetylcholine release from the frontal cortex during exploratory activity. Brain research 784, 218–227.

Glickman, S.E., and Sroges, R.W. (1966). Curiosity in zoo animals. Behaviour 26, 151–187.

Halassa, M.M., Florian, C., Fellin, T., Munoz, J.R., Lee, S.Y., Abel, T., Haydon, P.G., and Frank, M.G. (2009). Astrocytic modulation of sleep homeostasis and cognitive consequences of sleep loss. Neuron 61, 213–219. 10.1016/j.neuron.2008.11.024.

Hama, K., Arii, T., Katayama, E., Marton, M., and Ellisman, M.H. (2004). Tri-dimensional morphometric analysis of astrocytic processes with high voltage electron microscopy of thick Golgi preparations. Journal of neurocytology 33, 277–285.

Hasel, P., Dando, O., Jiwaji, Z., Baxter, P., Todd, A.C., Heron, S., Márkus, N.M., McQueen, J., Hampton, D.W., and Torvell, M. (2017). Neurons and neuronal activity control gene expression in astrocytes to regulate their development and metabolism. Nature communications 8, 1–18.

Haydon, P.G. (2017). Astrocytes and the modulation of sleep. Curr Opin Neurobiol 44, 28–33. 10.1016/j.conb.2017.02.008.

Hillman, E.M., Amoozegar, C.B., Wang, T., McCaslin, A.F., Bouchard, M.B., Mansfield, J., and Levenson, R.M. (2011). In vivo optical imaging and dynamic contrast methods for biomedical research. Philosophical Transactions of the Royal Society A: Mathematical, Physical and Engineering Sciences 369, 4620–4643.

Ho, W.S., and van den Pol, A.N. (2007). Bystander attenuation of neuronal and astrocyte intercellular communication by murine cytomegalovirus infection of glia. Journal of virology 81, 7286–7292.

Hrvatin, S., Hochbaum, D.R., Nagy, M.A., Cicconet, M., Robertson, K., Cheadle, L., Zilionis, R., Ratner, A., Borges-Monroy, R., and Klein, A.M. (2018). Single-cell analysis of experience-dependent transcriptomic states in the mouse visual cortex. Nature neuroscience 21, 120–129.

Hsu, J.-M., Kang, Y., Corty, M.M., Mathieson, D., Peters, O.M., and Freeman, M.R. (2021). Injury-induced inhibition of bystander neurons requires dSarm and signaling from glia. Neuron 109, 473–487. e475.

Ingiosi, A.M., and Frank, M.G. (2022). Noradrenergic Signaling in Astrocytes Influences Mammalian Sleep Homeostasis. Clocks & Sleep 4, 332–345.

Ingiosi, A.M., Hayworth, C.R., Harvey, D.O., Singletary, K.G., Rempe, M.J., Wisor, J.P., and Frank, M.G. (2020). A Role for Astroglial Calcium in Mammalian Sleep and Sleep Regulation. Curr Biol 30, 4373–4383.e4377. 10.1016/j.cub.2020.08.052.

Jhuang, H., Garrote, E., Yu, X., Khilnani, V., Poggio, T., Steele, A.D., and Serre, T. (2010). Automated home-cage behavioural phenotyping of mice. Nature communications 1, 1–10.

Kabra, M., Robie, A.A., Rivera-Alba, M., Branson, S., and Branson, K. (2013). JAABA: interactive machine learning for automatic annotation of animal behavior. Nature methods 10, 64–67.

Kaczmarczyk, L., Reichenbach, N., Blank, N., Jonson, M., Dittrich, L., Petzold, G.C., and Jackson, W.S. (2021). Slc1a3-2A-CreERT2 mice reveal unique features of Bergmann glia and augment a growing collection of Cre drivers and effectors in the 129S4 genetic background. Scientific reports 11, 1–18.

Katori, Y., Sakamoto, K., Saito, N., Tanji, J., Mushiake, H., and Aihara, K. (2011). Representational switching by dynamical reorganization of attractor structure in a network model of the prefrontal cortex. PLoS computational biology 7, e1002266.

Kellner, V., Kersbergen, C.J., Li, S., Babola, T.A., Saher, G., and Bergles, D.E. (2021). Dual metabotropic glutamate receptor signaling enables coordination of astrocyte and neuron activity in developing sensory domains. Neuron 109, 2545–2555. e2547.

Kemp, A.H., Gray, M.A., Silberstein, R.B., Armstrong, S.M., and Nathan, P.J. (2004). Augmentation of serotonin enhances pleasant and suppresses unpleasant cortical electrophysiological responses to visual emotional stimuli in humans. Neuroimage 22, 1084–1096.

Kjaerby, C., Andersen, M., Hauglund, N., Ding, F., Wang, W., Xu, Q., Deng, S., Kang, N., Peng, S., and Sun, Q. (2020). Dynamic fluctuations of the locus coeruleus-norepinephrine system underlie sleep state transitions. Biorxiv.

Kokare, D.M., Shelkar, G.P., Borkar, C.D., Nakhate, K.T., and Subhedar, N.K. (2011). A simple and inexpensive method to fabricate a cannula system for intracranial injections in rats and mice. Journal of pharmacological and toxicological methods 64, 246–250.

Kol, A., Adamsky, A., Groysman, M., Kreisel, T., London, M., and Goshen, I. (2020). Astrocytes contribute to remote memory formation by modulating hippocampal–cortical communication during learning. Nature neuroscience 23, 1229–1239.

Kozberg, M.G., Ma, Y., Shaik, M.A., Kim, S.H., and Hillman, E.M. (2016). Rapid postnatal expansion of neural networks occurs in an environment of altered neurovascular and neurometabolic coupling. Journal of Neuroscience 36, 6704–6717.

Lee, S., Meyer, J.F., Park, J., and Smirnakis, S.M. (2017). Visually driven neuropil activity and information encoding in mouse primary visual cortex. Frontiers in neural circuits 11, 50.

Lee, S.A., Holly, K.S., Voziyanov, V., Villalba, S.L., Tong, R., Grigsby, H.E., Glasscock, E., Szele, F.G., Vlachos, I., and Murray, T.A. (2016). Gradient index microlens implanted in prefrontal cortex of mouse does not affect behavioral test performance over time. PloS one 11, e0146533.

Lever, C., Burton, S., and Ο’Keefe, J. (2006). Rearing on hind legs, environmental novelty, and the hippocampal formation. Reviews in the Neurosciences 17, 111–134.

Lever, C., Wills, T., Cacucci, F., Burgess, N., and O’Keefe, J. (2002). Long-term plasticity in hippocampal place-cell representation of environmental geometry. Nature 416, 90–94.

Levy, S., and Bargmann, C.I. (2020). An adaptive-threshold mechanism for odor sensation and animal navigation. Neuron 105, 534–548. e513.

Lopes, G., Nogueira, J., Dimitriadis, G., Menendez, J.A., Paton, J.J., and Kampff, A.R. (2017). A robust role for motor cortex. bioRxiv, 058917. 10.1101/058917.

Lyon, K.A., and Allen, N.J. (2022). From Synapses to Circuits, Astrocytes Regulate Behavior. Frontiers in Neural Circuits, 136.

Ma, Y., Shaik, M.A., Kozberg, M.G., Kim, S.H., Portes, J.P., Timerman, D., and Hillman, E.M. (2016). Resting-state hemodynamics are spatiotemporally coupled to synchronized and symmetric neural activity in excitatory neurons. Proceedings of the National Academy of Sciences 113, E8463–E8471.

MacVicar, B.A. (1997). REVIEW▪: Mapping Neuronal Activity by Imaging Intrinsic Optical Signals. The Neuroscientist 3, 381–388.

Maloney, S.E., Tabachnick, D.R., Jakes, C., Avdagic, S., Bauernfeind, A.L., and Dougherty, J.D. (2022). Fluoxetine exposure throughout neurodevelopment differentially influences basilar dendritic morphology in the motor and prefrontal cortices. Scientific reports 12, 1–12.

Mariotti, L., Losi, G., Lia, A., Melone, M., Chiavegato, A., Gómez-Gonzalo, M., Sessolo, M., Bovetti, S., Forli, A., and Zonta, M. (2018). Interneuron-specific signaling evokes distinctive somatostatin-mediated responses in adult cortical astrocytes. Nature communications 9, 1–14.

Mederos, S., Hernández-Vivanco, A., Ramírez-Franco, J., Martín-Fernández, M., Navarrete, M., Yang, A., Boyden, E.S., and Perea, G. (2019). Melanopsin for precise optogenetic activation of astrocyte-neuron networks. Glia 67, 915–934.

Merten, K., Folk, R.W., Duarte, D., and Nimmerjahn, A. (2021). Astrocytes encode complex behaviorally relevant information. bioRxiv.

Miller, S.J., Philips, T., Kim, N., Dastgheyb, R., Chen, Z., Hsieh, Y.-C., Daigle, J.G., Datta, M., Chew, J., and Vidensky, S. (2019). Molecularly defined cortical astroglia subpopulation modulates neurons via secretion of Norrin. Nature neuroscience 22, 741–752.

Miranda, M.A.I., Ramıŕ ez-Lugo, L., and Bermúdez-Rattoni, F. (2000). Cortical cholinergic activity is related to the novelty of the stimulus. Brain research 882, 230–235.

Miyamoto, D., and Murayama, M. (2016). The fiber-optic imaging and manipulation of neural activity during animal behavior. Neuroscience Research 103, 1–9.

Moeyaert, B., Holt, G., Madangopal, R., Perez-Alvarez, A., Fearey, B.C., Trojanowski, N.F., Ledderose, J., Zolnik, T.A., Das, A., and Patel, D. (2018). Improved methods for marking active neuron populations. Nature communications 9, 1–12.

Montijn, J.S., Vinck, M., and Pennartz, C. (2014). Population coding in mouse visual cortex: response reliability and dissociability of stimulus tuning and noise correlation. Frontiers in computational neuroscience 8, 58.

Mu, Y., Bennett, D.V., Rubinov, M., Narayan, S., Yang, C.-T., Tanimoto, M., Mensh, B.D., Looger, L.L., and Ahrens, M.B. (2019). Glia accumulate evidence that actions are futile and suppress unsuccessful behavior. Cell 178, 27–43. e19.

Nagai, J., Bellafard, A., Qu, Z., Yu, X., Ollivier, M., Gangwani, M.R., Diaz-Castro, B., Coppola, G., Schumacher, S.M., and Golshani, P. (2021). Specific and behaviorally consequential astrocyte Gq GPCR signaling attenuation in vivo with iβARK. Neuron 109, 2256–2274. e2259.

Nandy, A.S., Nassi, J.J., and Reynolds, J.H. (2017). Laminar organization of attentional modulation in macaque visual area V4. Neuron 93, 235–246.

Navarrete, M., Perea, G., de Sevilla, D.F., Gómez-Gonzalo, M., Núñez, A., Martín, E.D., and Araque, A. (2012). Astrocytes mediate in vivo cholinergic-induced synaptic plasticity. PLoS biology 10, e1001259.

Nawrot, M.P. (2010). Analysis and interpretation of interval and count variability in neural spike trains. In Analysis of parallel spike trains, (Springer), pp. 37–58.

Nimmerjahn, A., and Bergles, D.E. (2015). Large-scale recording of astrocyte activity. Current opinion in neurobiology 32, 95–106.

Noguchi, K., Gel, Y.R., Brunner, E., and Konietschke, F. (2012). nparLD: an R software package for the nonparametric analysis of longitudinal data in factorial experiments. Journal of Statistical software 50, 1–23.

O’Haver, T. Pragmatic Introduction to Signal Processing 2019: Applications in scientific measurement. 2019.

Oberheim, N.A., Goldman, S.A., and Nedergaard, M. (2012). Heterogeneity of astrocytic form and function. Astrocytes, 23-45.

Oliveira, J.F., Sardinha, V.M., Guerra-Gomes, S., Araque, A., and Sousa, N. (2015). Do stars govern our actions? Astrocyte involvement in rodent behavior. Trends in neurosciences 38, 535–549.

Pak, R.W., Hsu, E., Molina-Castro, G., Bergles, D.E., and Kang, J.U. (2021). Fluorescence dual-color fiberscope for monitoring neuron and astrocyte concurrent activities in freely-behaving animals. (International Society for Optics and Photonics), pp. 116291N.

Papouin, T., Dunphy, J., Tolman, M., Foley, J.C., and Haydon, P.G. (2017a). Astrocytic control of synaptic function. Philosophical Transactions of the Royal Society B: Biological Sciences 372, 20160154.

Papouin, T., Dunphy, J.M., Tolman, M., Dineley, K.T., and Haydon, P.G. (2017b). Septal cholinergic neuromodulation tunes the astrocyte-dependent gating of hippocampal NMDA receptors to wakefulness. Neuron 94, 840–854. e847.

Paukert, M., Agarwal, A., Cha, J., Doze, V.A., Kang, J.U., and Bergles, D.E. (2014). Norepinephrine controls astroglial responsiveness to local circuit activity. Neuron 82, 1263–1270.

Pchitskaya, E., Popugaeva, E., and Bezprozvanny, I. (2018). Calcium signaling and molecular mechanisms underlying neurodegenerative diseases. Cell calcium 70, 87–94.

Peters, A.J., Lee, J., Hedrick, N.G., O’Neil, K., and Komiyama, T. (2017). Reorganization of corticospinal output during motor learning. Nature neuroscience 20, 1133–1141.

Plöschner, M., Kollárová, V., Dostál, Z., Nylk, J., Barton-Owen, T., Ferrier, D.E., Chmelík, R., Dholakia, K., and Čižmár, T. (2015). Multimode fibre: Light-sheet microscopy at the tip of a needle. Scientific reports 5, 1–7.

Poskanzer, K.E., and Yuste, R. (2016). Astrocytes regulate cortical state switching in vivo. Proceedings of the National Academy of Sciences 113, E2675–E2684.

Qin, H., He, W., Yang, C., Li, J., Jian, T., Liang, S., Chen, T., Feng, H., Chen, X., Liao, X., and Zhang, K. (2020). Monitoring Astrocytic Ca(2+) Activity in Freely Behaving Mice. Front Cell Neurosci 14, 603095. 10.3389/fncel.2020.603095.

Rector, D.M., Carter, K.M., Volegov, P.L., and George, J.S. (2005). Spatio-temporal mapping of rat whisker barrels with fast scattered light signals. Neuroimage 26, 619–627.

Refaeli, R., Doron, A., Benmelech-Chovav, A., Groysman, M., Kreisel, T., Loewenstein, Y., and Goshen, I. (2021). Features of hippocampal astrocytic domains and their spatial relation to excitatory and inhibitory neurons. Glia 69, 2378–2390.

Reitman, M.E., Tse, V., Mi, X., Willoughby, D.D., Peinado, A., Aivazidis, A., Myagmar, B.E., Simpson, P.C., Bayraktar, O.A., Yu, G., and Poskanzer, K.E. (2023). Norepinephrine links astrocytic activity to regulation of cortical state. Nat Neurosci. 10.1038/s41593-023-01284-w.

Requie, L.M., Gómez-Gonzalo, M., Speggiorin, M., Managò, F., Melone, M., Congiu, M., Chiavegato, A., Lia, A., Zonta, M., and Losi, G. (2022). Astrocytes mediate long-lasting synaptic regulation of ventral tegmental area dopamine neurons. Nature Neuroscience, 1–12.

Rungta, R.L., Bernier, L.P., Dissing-Olesen, L., Groten, C.J., LeDue, J.M., Ko, R., Drissler, S., and MacVicar, B.A. (2016). Ca2+ transients in astrocyte fine processes occur via Ca2+ influx in the adult mouse hippocampus. Glia 64, 2093–2103.

Sakers, K., Liu, Y., Llaci, L., Lee, S.M., Vasek, M.J., Rieger, M.A., Brophy, S., Tycksen, E., Lewis, R., and Maloney, S.E. (2021). Loss of Quaking RNA binding protein disrupts the expression of genes associated with astrocyte maturation in mouse brain. Nature communications 12, 1–14.

Salatino, J.W., Ludwig, K.A., Kozai, T.D., and Purcell, E.K. (2017). Glial responses to implanted electrodes in the brain. Nature biomedical engineering 1, 862–877.

Sapkota, D., Kater, M.S., Sakers, K., Nygaard, K.R., Liu, Y., Koester, S.K., Fass, S.B., Lake, A.M., Khazanchi, R., and Khankan, R.R. (2022). Activity-dependent translation dynamically alters the proteome of the perisynaptic astrocyte process. Cell reports 41, 111474.

Sardar, D., Lozzi, B., Woo, J., Huang, T.-W., Cvetkovic, C., Creighton, C.J., Krencik, R., and Deneen, B. (2021). Mapping astrocyte transcriptional signatures in response to neuroactive compounds. International journal of molecular sciences 22, 3975.

Schindelin, J., Arganda-Carreras, I., Frise, E., Kaynig, V., Longair, M., Pietzsch, T., Preibisch, S., Rueden, C., Saalfeld, S., and Schmid, B. (2012). Fiji: an open-source platform for biological-image analysis. Nature methods 9, 676–682.

Sekiguchi, K.J., Shekhtmeyster, P., Merten, K., Arena, A., Cook, D., Hoffman, E., Ngo, A., and Nimmerjahn, A. (2016). Imaging large-scale cellular activity in spinal cord of freely behaving mice. Nature communications 7, 1–13.

Sheintuch, L., Rubin, A., Brande-Eilat, N., Geva, N., Sadeh, N., Pinchasof, O., and Ziv, Y. (2017). Tracking the same neurons across multiple days in Ca2+ imaging data. Cell reports 21, 1102–1115.

Shekhtmeyster, P., Duarte, D., Carey, E.M., Ngo, A., Gao, G., Olmstead, J.A., Nelson, N.A., and Nimmerjahn, A. (2021). Trans-segmental imaging in the spinal cord of behaving mice. bioRxiv.

Sivankutty, S., Tsvirkun, V., Bouwmans, G., Kogan, D., Oron, D., Andresen, E.R., and Rigneault, H. (2016). Extended field-of-view in a lensless endoscope using an aperiodic multicore fiber. Optics letters 41, 3531–3534.

Slezak, M., Kandler, S., Van Veldhoven, P.P., Van den Haute, C., Bonin, V., and Holt, M.G. (2019). Distinct mechanisms for visual and motor-related astrocyte responses in mouse visual cortex. Current Biology 29, 3120–3127. e3125.

Soleimanzad, H., Smekens, F., Peyronnet, J., Juchaux, M., Lefebvre, O., Bouville, D., Magnan, C., Gurden, H., and Pain, F. (2019). Multiple speckle exposure imaging for the study of blood flow changes induced by functional activation of barrel cortex and olfactory bulb in mice. Neurophotonics 6, 015008.

Sołyga, M., and Barkat, T.R. (2019). Distinct processing of tone offset in two primary auditory cortices. Scientific reports 9, 1–12.

Song, C., Do, Q.B., Antic, S.D., and Knöpfel, T. (2017). Transgenic strategies for sparse but strong expression of genetically encoded voltage and calcium indicators. International Journal of Molecular Sciences 18, 1461.

Srinivasan, R., Huang, B.S., Venugopal, S., Johnston, A.D., Chai, H., Zeng, H., Golshani, P., and Khakh, B.S. (2015). Ca2+ signaling in astrocytes from Ip3r2−/− mice in brain slices and during startle responses in vivo. Nature neuroscience 18, 708–717.

Streich, L., Boffi, J.C., Wang, L., Alhalaseh, K., Barbieri, M., Rehm, R., Deivasigamani, S., Gross, C.T., Agarwal, A., and Prevedel, R. (2021). High-resolution structural and functional deep brain imaging using adaptive optics three-photon microscopy. Nature methods 18, 1253–1258.

Sweeney, P., Qi, Y., Xu, Z., and Yang, Y. (2016). Activation of hypothalamic astrocytes suppresses feeding without altering emotional states. Glia 64, 2263–2273.

Szabo, V., Ventalon, C., De Sars, V., Bradley, J., and Emiliani, V. (2014). Spatially selective holographic photoactivation and functional fluorescence imaging in freely behaving mice with a fiberscope. Neuron 84, 1157–1169.

Szabó, Z., Héja, L., Szalay, G., Kékesi, O., Füredi, A., Szebényi, K., Dobolyi, Á., Orbán, T.I., Kolacsek, O., and Tompa, T. (2017). Extensive astrocyte synchronization advances neuronal coupling in slow wave activity in vivo. Scientific reports 7, 1–18.

Takata, N., Mishima, T., Hisatsune, C., Nagai, T., Ebisui, E., Mikoshiba, K., and Hirase, H. (2011). Astrocyte calcium signaling transforms cholinergic modulation to cortical plasticity in vivo. Journal of Neuroscience 31, 18155–18165.

Thrane, A.S., Rangroo Thrane, V., Zeppenfeld, D., Lou, N., Xu, Q., Nagelhus, E.A., and Nedergaard, M. (2012). General anesthesia selectively disrupts astrocyte calcium signaling in the awake mouse cortex. Proc Natl Acad Sci U S A 109, 18974–18979. 10.1073/pnas.1209448109.

Toates, F.M. (1986). Motivational systems (CUP Archive).

Tsunematsu, T., Sakata, S., Sanagi, T., Tanaka, K.F., and Matsui, K. (2021). Region-Specific and State-Dependent Astrocyte Ca(2+) Dynamics during the Sleep-Wake Cycle in Mice. J Neurosci 41, 5440–5452. 10.1523/jneurosci.2912-20.2021.

Ung, K., Tepe, B., Pekarek, B., Arenkiel, B.R., and Deneen, B. (2020). Parallel astrocyte calcium signaling modulates olfactory bulb responses. Journal of neuroscience research 98, 1605–1618.

Vaidyanathan, T.V., Collard, M., Yokoyama, S., Reitman, M.E., and Poskanzer, K.E. (2021). Cortical astrocytes independently regulate sleep depth and duration via separate GPCR pathways. Elife 10. 10.7554/eLife.63329.

Vecoven, N., Ernst, D., Wehenkel, A., and Drion, G. (2020). Introducing neuromodulation in deep neural networks to learn adaptive behaviours. PloS one 15, e0227922.

Welker, W. (1957). “Free” versus “forced” exploration of a novel situation by rats. Psychological Reports 3, 95–108.

Xie, Y., Kuan, A.T., Wang, W., Herbert, Z.T., Mosto, O., Olukoya, O., Adam, M., Vu, S., Kim, M., and Tran, D. (2022). Astrocyte-neuron crosstalk through Hedgehog signaling mediates cortical synapse development. Cell reports 38, 110416.

Xu, L., Anwyl, R., and Rowan, M.J. (1998). Spatial exploration induces a persistent reversal of long-term potentiation in rat hippocampus. Nature 394, 891–894.

Yang, L., Qi, Y., and Yang, Y. (2015). Astrocytes control food intake by inhibiting AGRP neuron activity via adenosine A1 receptors. Cell reports 11, 798–807.

Yu, M., Li, Y., and Qiu, M. (2019). Statistical inference of the relative concentration index for complex surveys. Statistics in Medicine 38, 4083–4095.

Yu, X., Moye, S.L., and Khakh, B.S. (2021). Local and CNS-wide astrocyte intracellular calcium signaling attenuation in vivo with CalExflox mice. Journal of Neuroscience 41, 4556–4574.

Yu, X., Nagai, J., Marti-Solano, M., Soto, J.S., Coppola, G., Babu, M.M., and Khakh, B.S. (2020). Context-specific striatal astrocyte molecular responses are phenotypically exploitable. Neuron 108, 1146–1162. e1110.

Zamanian, J.L., Xu, L., Foo, L.C., Nouri, N., Zhou, L., Giffard, R.G., and Barres, B.A. (2012). Genomic analysis of reactive astrogliosis. Journal of neuroscience 32, 6391–6410.

Zhang, K., Förster, R., He, W., Liao, X., Li, J., Yang, C., Qin, H., Wang, M., Ding, R., Li, R., et al. (2021). Fear learning induces α7-nicotinic acetylcholine receptor-mediated astrocytic responsiveness that is required for memory persistence. Nat Neurosci 24, 1686–1698. 10.1038/s41593-021-00949-8.

Zhou, P., Resendez, S.L., Rodriguez-Romaguera, J., Jimenez, J.C., Neufeld, S.Q., Giovannucci, A., Friedrich, J., Pnevmatikakis, E.A., Stuber, G.D., and Hen, R. (2018). Efficient and accurate extraction of in vivo calcium signals from microendoscopic video data. Elife 7, e28728.

